# A semi-automated genetic screen using genome-wide association analysis identifies modulators of stress-induced sleep in *Caenorhabditis elegans*

**DOI:** 10.64898/2026.05.16.725661

**Authors:** Zihao (John) Li, Helya Honarpisheh, Sreekar Kutagulla, Kerry M. LeCure, Jing Liang, David M. Raizen, Christopher Fang-Yen

## Abstract

Animals sleep more when they are sick. In *C. elegans*, stress-induced sleep (SIS) follows cellular injury such as exposure to ultraviolet (UV) light. The genetic regulators of SIS remain incompletely defined. Using robotic automation, high-throughput imaging, and genome-wide association analysis, we performed a semi-automated screen of 940 whole-genome sequenced Million Mutation Project (MMP) strains. We quantified behavioral activity and quiescence before and after ultraviolet (UV) radiation. We applied the Sequence Kernel Association Test (SKAT) to this behavioral data to prioritize a set of associated genes and observed significant enrichment of known SIS genetic regulators. Based on these results, we conducted a candidate validation screen for additional genes regulating SIS. We identified three genes (*strd-1, cla-1,* and *egl-8),* mutations in which reproducibly influence SIS. Further exploration of these genes holds potential for enhancing our understanding of the molecular basis of SIS. These findings establish a pipeline for automated behavioral phenotyping coupled with gene-based association to accelerate studies of *C. elegans* neurogenetics.

## Introduction

Sleep is a fundamental yet poorly understood biological process. Many animals enter a sleep or fatigue state during periods of sickness (Besedovsky et al., 2019; Dantzer et al., 2008; Lopes et al., 2021). In response to infection, cellular stress, or tissue injury, animals exhibit a collection of adaptive behaviors, including increased sleepiness, reduced foraging, inhibited movement, and withdrawal from social engagement (Davis & Raizen, 2017). In invertebrates, stress induced sleep (SIS) is characterized by a cessation or near cessation of locomotion and feeding and an increased threshold to arousal by sensory stimuli (Hill et al., 2014; Trojanowski & Raizen, 2016). Environmental factors that cause SIS in fruit flies and roundworms include temperature shock, osmotic shock, toxins, pathogens, and exposure to ultraviolet light (DeBardeleben et al., 2017; Hill et al., 2014; Iannacone et al., 2024; Lenz et al., 2015; Pradhan et al., 2025). SIS is beneficial for animals to recover from injuries. In both *C. elegans* (Fry et al., 2016; Hill et al., 2014) and *Drosophila* (Kuo & Williams, 2014; Lenz et al., 2015), animals with defective SIS show elevated mortality following exposure to stress.

Prior work has described some of the molecular mechanisms underlying *C. elegans* SIS. After cellular stress, Epidermal Growth Factor (EGF) ligands LIN-3 (Hill et al., 2014) and SISS-1 (Hill et al., 2024) target the sleep-promoting interneurons ALA (Van Buskirk & Sternberg, 2007) and RIS (Konietzka et al., 2020), which then release neuropeptides to trigger sleep. KIN-29, the *C. elegans* ortholog of salt-inducible kinase (SIK), is another important SIS regulator that acts upstream of ALA and RIS (Grubbs et al., 2020). KIN-29 functions in sensory neurons that respond to metabolic signals under stress, promoting activation of ALA and RIS to induce sleep (Grubbs et al., 2020).

Downstream of ALA and RIS interneurons, neuropeptide signaling plays an important role in sleep induction. Neuropeptides including FLP-13 (Nelson et al., 2014), FLP-24, and NLP-8 (Nath et al., 2016) downstream of ALA, and FLP-11 downstream of RIS (Busack & Bringmann, 2023), inhibit wake-promoting neurons to induce sleep. Recent work has shown that the ionotropic glutamate receptor GLR-5 also contributes to SIS by modulating glutamate signaling within RIS circuitry (Howe et al., 2026).

Although these important components are known, the broader genetic architecture of SIS remains unclear. Forward genetic screening is a powerful approach for deciphering the genetic mechanisms underlying animals’ sleep. Genetic screens for sleep-regulating candidates have been conducted in worms (Huang et al., 2017a; Iannacone et al., 2017; Sinner et al., 2021), flies (Cirelli et al., 2005; Toda et al., 2019; Wu et al., 2008), fish (Barlow et al., 2023; Chiu et al., 2016), and mice (Funato et al., 2016; Keenan et al., 2021).

Genetics screens in *C. elegans* are typically performed using manual methods, which are labor intensive and low in throughput. A prior screen discovered a single allele of the gene *aptf-1* through manually looking for populations without any immobile larvae (Turek et al., 2013). Another non-clonal screen identified the gene *npr-1* as necessary for SIS in *C. elegans* (Soto et al., 2019). Due to the limited throughput, these screens identified only a few alleles for the single gene and were far from saturation. While screens for suppressors of the gene *flp-13* (Iannacone et al., 2017; Yuan et al., 2015) were performed in a relatively high-throughput manner and probably reached saturation, it focused only on a particular aspect of sleep regulation (downstream of *flp-13*).

The central goal of genetic screens is to identify genes associated with a phenotype of interest. However, a major bottleneck in classical forward genetic screens is to identify and map the genes with moderate effects, because these approaches often prioritize mutants with strong phenotypes. The Million Mutation Project (MMP) provides a library of 2,007 mutagenized, whole-genome-sequenced strains (Thompson et al., 2013), enabling phenotypes to be associated with genetic variants across the genome. The Sequence Kernel Association Test (SKAT) is a regression-based method that tests the combined association of variants within a gene or genomic region with a continuous or discrete phenotype (Wu et al., 2011). Applied to MMP strains, SKAT can prioritize genes associated with phenotypic variation, including genes with moderate effects that might be difficult to identify in classical genetic screens. This strategy was previously used to identify genes associated with dye-filling defects in *C. elegans* ciliated sensory neurons (Timbers et al., 2016). However, to our knowledge, it has not previously been applied to a *C. elegans* behavioral screen.

In this study, we conducted a genetic screen for regulators of UV-induced sleep using strains from the MMP library (Thompson et al., 2013). The screen required long-term behavioral assays for large numbers of animals, making the experiments labor intensive. To increase throughput, we integrated automation technologies into the screening workflow, including high-throughput imaging using the WorMotel platform (Churgin et al., 2017) and automated *C. elegans* picking using the WormPicker robot (Li et al., 2023). We used WormPicker (Li et al., 2023) to maintain the MMP strain library and prepare sleep assays in the multi-well WorMotel platform (Churgin et al., 2017). Using a combination of manual and automated methods, we assayed the behavioral quiescence of adult animals before and after UV exposure across 940 strains randomly selected from the MMP library.

Using the sleep data from these MMP strains and their genomic sequence, we applied SKAT (Timbers et al., 2016; Wu et al., 2011) to identify genes statistically associated with variation in the SIS phenotype. We generated a ranking for 6,663 genes for which at least five non-synonymous alleles were represented in our collection of mutagenized strains, based on their statistical significance in the SKAT analysis. This ranking highlighted multiple genes already known to regulate SIS and identified novel candidates. We conducted a candidate screen based on this ranking and identified three genes, *strd-1*, *cla-1*, and *egl-8*, mutations in which reproducibly influence SIS. Notably, these genes are associated with moderate changes in the sleep phenotype that might be difficult to map using conventional genetic methods, demonstrating the advantages of our genome-wide association approach.

## Materials and methods

### Design of a semi-automated screen of SIS

We designed a pipeline for a semi-automated genetic screen for SIS regulators (Fig. 1A) using a library of mutagenized and sequenced strains from the MMP (Thompson et al., 2013). To reduce the labor involved in strain maintenance, we used our recently developed worm picking robot (Li et al., 2023) to perform timed egg laying to generate synchronized populations for sleep assays of individual MMP strains. For each MMP strain, the robot picked 5-10 gravid adults to fresh plates, allowed them to lay embryos for 4 hours, then removed the worms. To ensure that only embryos remained on the plates, the automated system was able to eliminate more than 99% (N = 113 animals) of the gravid animals previously transferred. Experiments in which the adults were not fully eliminated were censored and the trials were repeated using the robot. For some strains (∼10%) that were egg laying defective, the robot was used to pick L4 animals one day before SIS phenotyping.

**Fig. 1.**
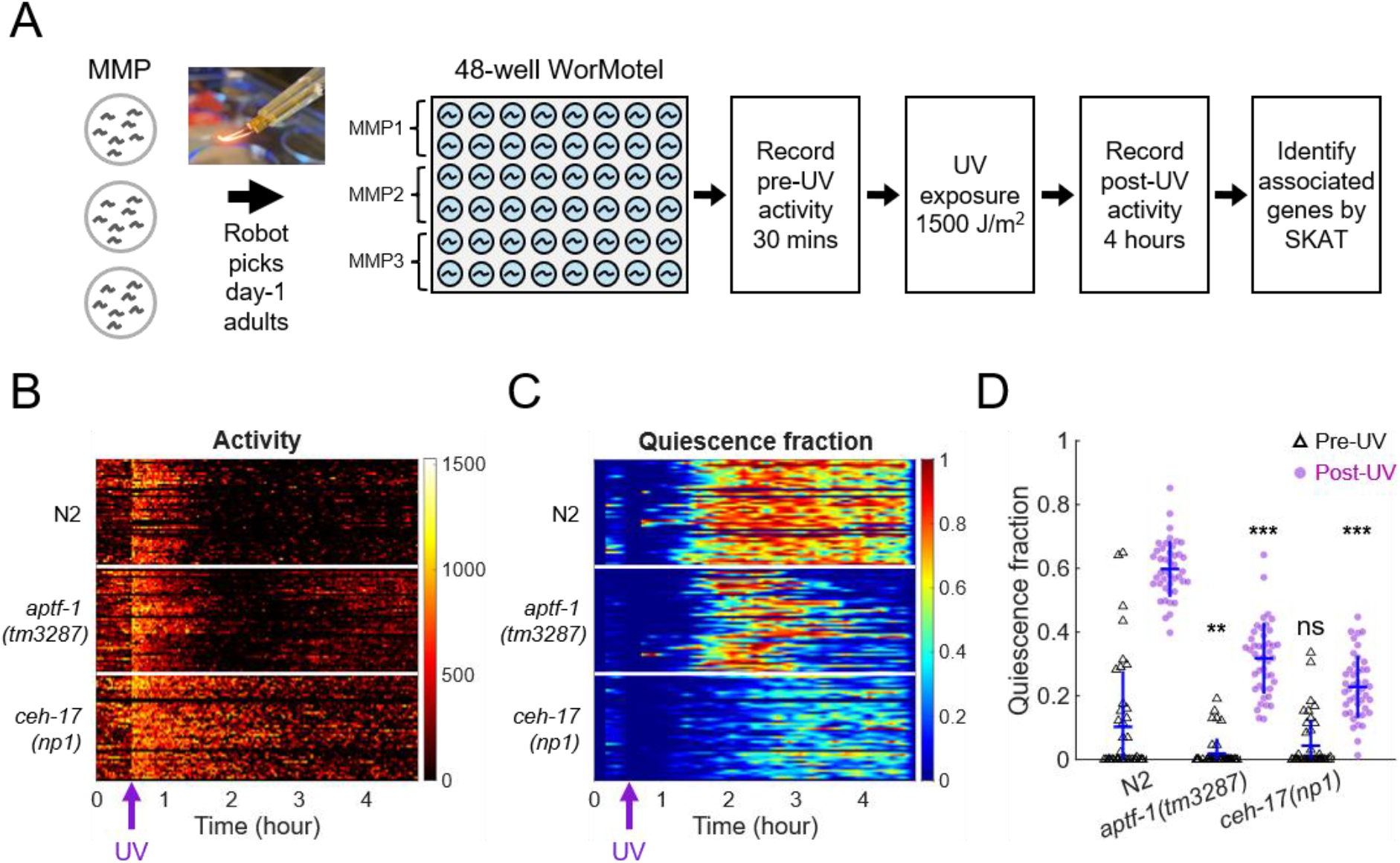
Design of automated genetic screen for modifiers of SIS in *C. elegans*. (A) Schematic diagram of the workflow. The photo of an electrically-sterilized robotic worm pick is adapted from Li et al. (2023). (B and C) Time course of pre-UV and post-UV (B) activity and (C) quiescence for individual animals of N2 and SIS mutants *aptf-1(tm3287)* and *ceh-17(np1)*. Each row represents one animal. (D) Total pre-UV and post-UV quiescence fraction for N2, *aptf-1(tm3287)*, and *ceh-17(np1)*. Each point represents a single animal. The error bars represent one standard deviation above and below the mean. Statistics were performed using one-sided Wilcoxon rank-sum tests, examining whether the pre- and post-UV quiescence fractions of each mutant are lower than those of N2. \*\*\**p* < 0.001 and \*\**p* < 0.01, ns: not significant. For (B)-(D), N = 43, 44, and 44 animals for N2, *aptf-1(tm3287)*, and *ceh-17(np1)*, respectively.

Due to the large inter-individual variability in *C. elegans* sleep (Raizen et al., 2008), reliable assessment requires a population of animals for each strain. To assay SIS, we used a microfabricated multi-well PDMS chip (WorMotel) optimized for longitudinal imaging of individual animals in each well (Churgin et al., 2017). For each 48-well WorMotel, the robotic system transferred three MMP strains (16 animals per strain) to individual wells on the chip (Fig. 1A and Movie S1). One group of N2 animals was included as a control for approximately every 30 MMP strains.

In addition to the automated workflow, some of the MMP strains were screened manually (Table S1). For these strains, the protocols for strain preparation and assay setup were identical to those used in the robotic workflow, except that the animals were picked manually. The throughput of the robot-automated screen (∼30 strains per week) was ∼50% higher than pure manual methods (∼20 strains per week).

The loaded WorMotel chips, prepared using robotic or manual methods, were transferred to imaging systems (Churgin & Fang-Yen, 2022) that recorded images for at least 30 minutes before UV treatment (1,500 J/m^2^ dosage) and for 4 hours after UV treatment (Fig. 1A), at a rate of one frame every 10 seconds. The level of quiescence was evaluated using the pixel difference obtained by subtracting every pair of temporally-adjacent frames (Churgin et al., 2019) and measuring how many pixel values changed beyond a threshold value. We summed the level of quiescence over the imaging period to determine the total amount of quiescence for each strain.

To validate our methods, we first used the robot to assay the behavioral activity and quiescence for the N2 control and two previously described sleep-defective mutants, *aptf-1* (Turek et al., 2013) and *ceh-17* (Hill et al., 2014) (Fig. 1B-D). Before the UV exposure, *aptf-1(tm3287)* mutants showed lower quiescence than N2, while *ceh-17(np1)* mutants showed comparable quiescence as N2. After UV exposure, both mutants showed significantly less quiescence than N2. These results show that the robotic method generates results for SIS consistent with published data.

### Preparation of *C. elegans* strains

*C. elegans* was cultivated on nematode growth medium (NGM) plates with OP50 bacteria at 20°C using standard methods (Stiernagle, 2021). Details of strains are given in Table S1. All experiments were carried out at ambient temperature (20 ± 1 °C).

### Robotic *C. elegans* picking system

An automated worm picking robot was constructed as described in Li et al. (2023). We developed custom machine vision methods to enable the robot to recognize individual wells on the WorMotel. The algorithm first identifies four corners of the WorMotel in the field of view (FOV) using template matching methods as described in the OpenCV library (Bradski, 2000). Next, coordinates of individual wells are inferred based on the geometrical parameters defined by the WorMotel geometry (Churgin et al., 2019).

### WorMotel multi-well substrates

The WorMotel is a microfabricated PDMS substrate containing an array of wells for longitudinal imaging of a population of *C. elegans* confined in individual wells (Churgin et al., 2017; Churgin et al., 2019). Each well of a 48-well WorMotel was filled with 15 µL NGM agar and 3.5 µL deionized water was pipetted on top of the agar in each well to facilitate worm loading. Next, animals grown on OP50-seeded agar plates were individually transferred into each well, along with a small amount of bacterial food, using either the worm picking robot or manual picking. After the water dried out or was absorbed by the agar, the WorMotel was placed in a 10 cm petri dish with the lid on and then transferred to an imaging station (Churgin & Fang-Yen, 2022) for behavioral phenotyping.

To prevent the agar from drying out in the wells during the imaging session, humidity was maintained by placing a moistened piece of lab tissue paper rolled up along the rim of the 10 cm petri dish (Churgin et al., 2019). To minimize water condensation on the lid, a thin layer of 20% Tween 20 solution was applied to the inner surface of the lid (Churgin et al., 2019).

### UV treatment to induce sleep in *C. elegans*

We induced sleep in adult *C. elegans* by exposing animals on WorMotel substrates to UV light (254 nm wavelength; 1,500 J/m^2^ dosage) delivered by a UV crosslinker (Spectrolinker XL-1000, Spectronics Corporation).

### Automated imaging of sleep

We set up imaging systems to monitor animals on the WorMotel, following the procedures described in Churgin and Fang-Yen (2022). Each WorMotel was imaged using dark\ field illumination created by red LED strips placed underneath a transparent glass stage. A camera (Imaging Source DMK 33GP031) equipped with a machine vision lens (Fujinon CF25HA-1, 25 mm focal length) was positioned above the WorMotel. The system was enclosed with black curtains to block ambient light. The camera captured one frame (2592 pixels x 1944 pixels, 4 cm x 3 cm FOV) every 10 seconds. Animals on the WorMotel were imaged for 30 minutes before and 4 hours after the UV treatment.

### Measurements of activity and quiescence

The methods reported in Churgin et al. (2019) were used to quantify animals’ activities and quiescence. To measure activity, two consecutive frames taken 10 seconds apart were compared, and the proportional change in each pixel (Δ*p*) was calculated using:

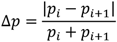

Here, *p_i_* represents the pixel gray scale value in the *i*-th frame and *p_i_*_+1_ represents the pixel gray scale value in the *i*+1-th frame. We defined activity as the number of pixels for which Δ*p* > 0.6.

In order to set an activity threshold to quantify quiescence, it is important to account for the baseline spontaneous activity for each strain, because genetic mutations can result in lower or higher overall locomotor activity (Schwarz & Bringmann, 2013; Yemini et al., 2013). Therefore, for each imaging trial, we first estimated the maximum activity level (MAL) for individual strains by calculating the median of the 95^th^ percentile of the activity recorded over the 30 mins before UV treatment:

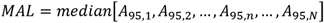

Here, *A*_95,*n*_ represents the 95^th^ percentile of the activity recorded over the 30 minutes before the UV treatment for the *n*-th animal, and *N* is the total number of animals in the imaging trial (*N* = 8 to 30 animals). We considered an animal to be quiescent at time *t* if the activity at that time is less than one percent of *MAL*.

We calculated the quiescence time over an imaging session as the total time during which the activity is less than one percent of *MAL*. The quiescence fraction is defined as the ratio between the total quiescence time and the total imaging time.

### SKAT analysis

We performed the analysis using the SKAT package (version 1.0.9) (Wu et al., 2011) in R (version 2023.12.0+369). First, genomic data from the MMP (Thompson et al., 2013) was used to construct a binary strain-genotype matrix ***G*** indicating which of the annotated genes contains non-synonymous mutations. Next, the SKAT model can be written as *S_i_* = *α*_0_ + ***G****_i_****β*** + *ε_i_*, where *S_i_* is the post-UV mean quiescence for the *i*-th MMP strain carrying non-synonymous mutations in the genes ***G****_i_*, *α*_0_ the intercept term, and *ε_i_* is the error term. No covariates were included in the model (Timbers et al., 2016). Under the null hypothesis of *β* = 0, we calculated *p* values for individual genes, followed by a Bonferroni correction. Smaller *p* values are considered as stronger association with the SIS phenotype. By sorting the *p* values, we generated a ranked list for the 18,070 genes, covering more than 90% of the *C. elegans* genome. On average, 4.64 ± 4.65 (mean ± SD) non-synonymous alleles were tested for each gene in our screen.

### Statistics

We performed statistical analyses using MATLAB R2025b. All two-sample comparisons of medians and variances were conducted using Wilcoxon rank-sum tests and Brown-Forsythe tests, respectively. Statistical details are provided in figures and their captions.

The enrichment of known sleep-regulating genes in the SKAT analysis was tested using randomization inference with N = 10,000 randomization trials under the null hypothesis that the known genes are randomly distributed throughout the ranked list.

## Results

### Semi-automated screen for SIS-modulating genes

Using a combination of automated and manual methods (Materials and methods), we assayed SIS phenotypes for 940 MMP strains (Fig. 2). Of these, 189 were obtained using the robot and 751 by manual methods (Table S1 and Discussion). In Fig. 2A, we show the distribution of quiescence fraction for 30 minutes before and 4 hours after the UV treatment for all the MMP strains we tested. Median quiescence of the MMP collection was higher than N2 for both before and after the UV treatment (Fig. 2A). We also observed that the variances of pre-UV and post-UV quiescence of the MMP strains are higher than these parameters in N2 (Fig. 2A). For most of the MMP strains, behavioral quiescence increased after exposure to UV (Fig. 2B).

**Fig. 2.**
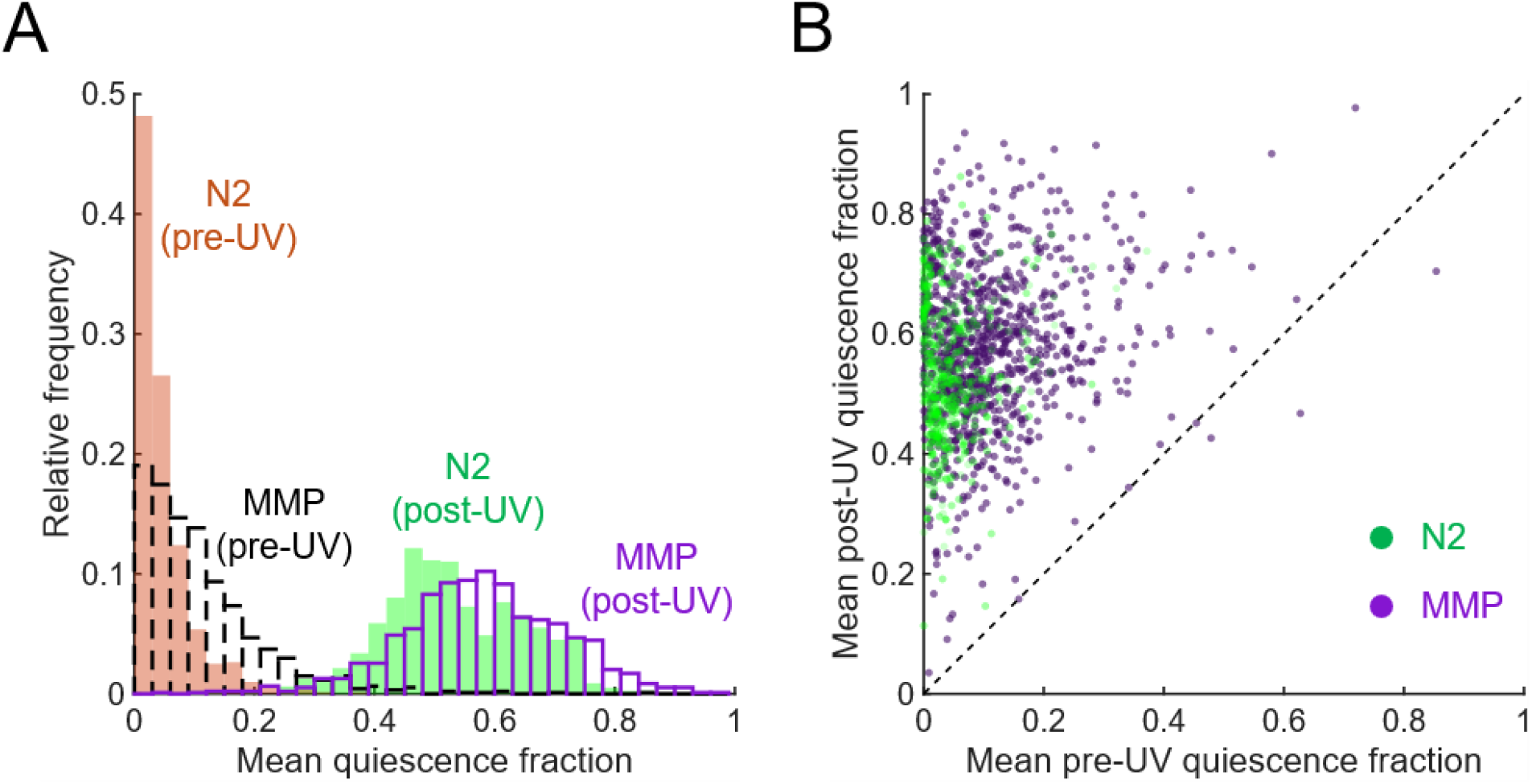
Screen for SIS phenotypes in MMP strains. (A) Distributions of mean pre-UV and post-UV quiescence fraction for N2 and the screened MMP strains. The median quiescence fractions of the MMP strains were compared with those of N2 using one-sided Wilcoxon rank-sum tests (pre-UV, ***; post-UV, ***). Equality of variances in quiescence fractions between the MMP strains and N2 was tested using Brown-Forsythe tests (pre-UV, ***; post-UV, **). \*\*\**p* < 0.001 and \*\**p* < 0.01. (B) A scatter plot showing the mean post-UV quiescence fraction versus pre-UV for N2 and the individual MMP strain. Each dot represents a single imaging trial for N2 or an individual MMP strain. The dashed line indicates where pre-UV and post-UV quiescence fraction would be equal. For (A) and (B), N = 799 imaging trials for N2; N = 940 MMP strains; N = 1 to 7 imaging trials for individual MMP strains; N = 8 to 30 animals for individual imaging trials; and N = 8 to 118 animals in total for individual MMP strains.

### SKAT identified candidate genes associated with altered SIS

Combining the SIS data and genomic data for the MMP strains, we used a linear regression-based method, SKAT (Timbers et al., 2016; Wu et al., 2011), to identify genes statistically associated with the SIS phenotype (Materials and methods). Based on the statistical significance of association revealed by SKAT, we generated a ranked list for 18,070 genes (Materials and methods).

As an initial verification of our SKAT analysis, we ask to what extent our gene list is able to identify known sleep-regulating genes in *C. elegans* from the literature (Tables 1 and S2). The list of known genes includes *ceh-17* (Hill et al., 2014), *ceh-14* (Van Buskirk & Sternberg, 2010), *aptf-1* (Turek et al., 2013), *dmsr-1* (Iannacone et al., 2017), *hda-4* (Grubbs et al., 2020), *siss-1* (Hill et al., 2024), *unc-108* (Robinson & Van Buskirk, 2019), *goa-1* (Huang et al., 2017b), *npr-1* (Soto et al., 2019), *npr-4* (Turek et al., 2016), *flp-11* (Turek et al., 2016), *grk-2* (Davis et al., 2023), *eel-1* (Raizen Lab, unpublished), and *rom-4* (Raizen Lab, unpublished). We also identified *eat-6*, a *C. elegans* Na-K-ATPase α-subunit (Davis et al., 1995) that regulates sleep in zebrafish (Barlow et al., 2023). We observed that the known genes clustered in the upper portion of the ranked list, with a median of rank percentile (MRP) of 17.9% (Tables 1 and S2). Under the null hypothesis that the known genes are randomly distributed throughout the ranked gene list, with an expected MRP of 50%, we obtain a *p* value of 0.002 (N = 10,000 randomization trials), supporting the idea that our SKAT analysis enriches for genes regulating SIS.

To further optimize the predictive power and confidence of our SKAT model, we filtered out genes represented by too few alleles by setting a threshold (*N_a_*) for the minimal number of non-synonymous alleles tested. We optimized the value of *N_a_* by minimizing the MRP for the known genes (Tables 1 and S2) and obtained an optimized value for *N_a_* = 5. With *N_a_* = 5, all the known genes cluster within the top 25% in our list. Based on these results, we reasoned that there may be other novel SIS regulators with high ranking in our list.

We acknowledge that the known genes listed above do not constitute a complete set of sleep-regulating genes. During peer review of this work, following a reviewer’s suggestion, we expanded the list by adding other known genes, including *let-23* (Hill et al., 2014; Van Buskirk & Sternberg, 2007), *kin-29* (Grubbs et al., 2020), *adm-4* (Hill et al., 2024), *sta-2* (Sinner et al., 2021), and *frm-10* (Raizen Lab, unpublished) (Table S2). We found that the test for enrichment of known genes remains statistically significant (*p* = 0.001, N = 10,000 randomization trials). The trend in MRP at different *N_a_* is similar to that shown in Table 1, reaching a minimum at *N_a_* = 3, indicating that the optimization method remains valid. Because the downstream candidate screen has been conducted based on the ranked list optimized using the initial set of known genes (*N_a_* = 5), we present the analysis results for the initial gene set in Table 1 and those for the expanded gene set in Table S2. The full lists of ranked genes for *N_a_* = 5 and *N_a_* = 3 are available in Tables S3 and S4, respectively.

**Table 1.** Effect of the threshold (*N_a_*) for the minimal counts of non-synonymous alleles in the SKAT analysis.

| $N_a$ | 1 | 2 | 3 | 4 | 5 | 6 | 7 | 8 | 9 |
| --- | --- | --- | --- | --- | --- | --- | --- | --- | --- |
| MRP* | 17.9 | 18.2 | 12.8 | 13.4 | 12.3 | 17.3 | 20.8 | 20.9 | 21.3 |
| $N_{kg}^{\dagger}$ | 15 | 13 | 9 | 7 | 6 | 4 | 3 | 3 | 3 |
| $N_{total}^{\ddagger}$ | 18,070 | 14,903 | 11,737 | 8,836 | 6,663 | 4,959 | 3,745 | 2,798 | 2,162 |

\*MRP denotes the median of rank percentile for the known sleep-regulating genes with the non-synonymous allele count not less than $N_a$ .
$^{\dagger}N_{kg}$ denotes the number of known sleep-regulating genes with the non-synonymous allele count greater than or equal to $N_a$ .
$^{\ddagger}N_{total}$ denotes the total number of genes with the non-synonymous allele count greater than or equal to $N_a$ .

### Candidate screen for genetic regulators of SIS

Equipped with the list of the top candidate genes generated by our SKAT analysis, we sought to experimentally identify genetic regulators for SIS. Based on the availability of strains in the lab’s frozen stock and from public genetic resources, including the NIH-funded *Caenorhabditis* Genetics Center and the National BioResource Project (Japan), we curated a list of strains with mutations in the high-ranking candidate genes (Table S1). We screened a total of 97 mutant strains for 67 top candidate genes (Fig. 3A and Table S1). For both before and after the UV exposure, the median quiescence for the mutant strains was higher than for N2. The variance of pre-UV quiescence for the mutant strains was not significantly different than that of N2. In contrast, after the exposure to UV, the mutant strains show larger variation in quiescence than N2. A gene was identified as a hit if at least two independent mutant alleles show consistent change in SIS compared to the N2 simultaneously imaged. When identifying mutants showing increased SIS, we required that they show pre-UV quiescence that was not significantly higher than that of N2, but post-UV quiescence that was higher than that of N2. This criterion reduces the likelihood that a mutant shows higher post-UV quiescence due to higher spontaneous quiescence. Our candidate screen identified three genes regulating SIS: *strd-1*, *cla-1*, and *egl-8* (Fig. 3B).

**Fig. 3.**
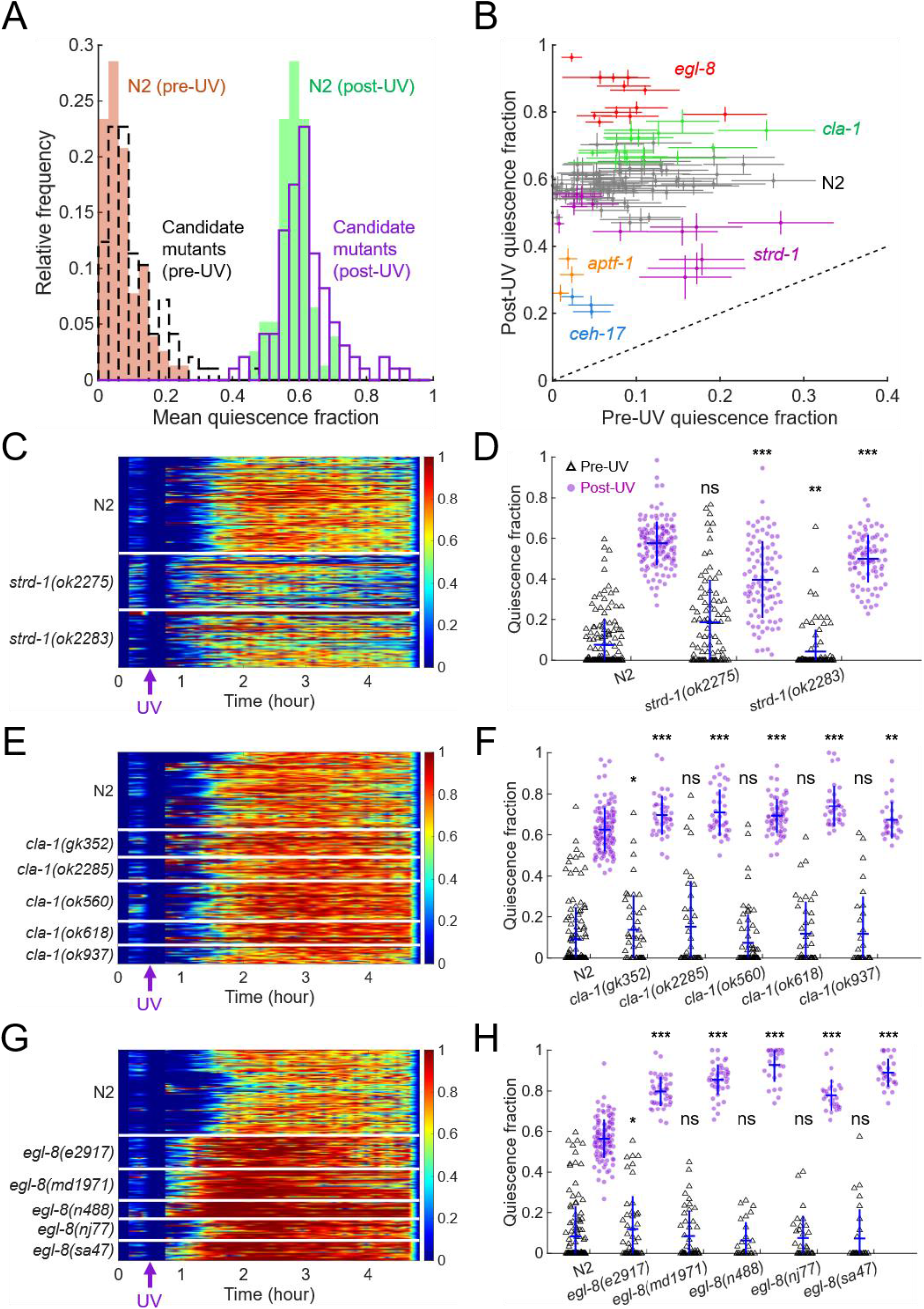
A candidate screen revealed novel genetic modulators of SIS. (A) Distribution of mean pre-UV and post-UV quiescence fraction for N2 (N = 77 imaging trials) and candidate mutants (N = 97 strains). The median quiescence fractions of the mutant strains were compared with those of N2 using one-sided Wilcoxon rank-sum tests (pre-UV, **; post-UV, **). Equality of variances in quiescence fractions between the mutant strains and N2 was tested using Brown-Forsythe tests (pre-UV, ns; post-UV, **). (B) Post-UV quiescence fraction versus pre-UV quiescence fraction for N2 and the SIS-altered mutants (N = 125 imaging trials). Individual points represent the mean across N = 6 to 24 animals for individual imaging trials. Error bars represent mean ± standard error. Dashed line represents where post-UV and pre-UV quiescence fraction would be equal. (C-H) Time course of quiescence (C, E, and G) and total quiescence fraction (D, F, and H) for independent alleles of (C and D) *strd-1*, (E and F) *cla-1*, and (G and H) *egl-8*, and N2 controls imaged simultaneously. Each horizontal row in the heat maps represents one animal. In the scatter plots, each dot represents one animal and error bars show one standard deviation above and below the mean. The legend in (D) applies to (F) and (H). For (C and D), N = 151 animals for N2, 89 for *strd-1(ok2275)*, 90 for *strd-1(ok2283)*. For (E and F), N = 128 animals for N2, 45 for *cla-1(gk352)*, 39 for *cla-1(ok2285)*, 67 for *cla-1(ok560)*, 39 for *cla-1(ok618)*, and 31 for *cla-1(ok937)*. For (G and H), N = 123 animals for N2, 48 for *egl-8(e2917)*, 45 for *egl-8(md1971)*, 27 for *egl-8(n488)*, 31 for *egl-8(nj77)*, 29 for *egl-8(sa47)*. For (D), (F), and (H), statistics were performed using one-sided Wilcoxon rank-sum tests, examining whether the pre- and post-UV quiescence fractions of each mutant are lower (D) or higher (F and H) than that of N2. \*\*\**p* < 0.001, \*\**p* < 0.01, and \**p* < 0.05, ns: not significant.

### Mutants for STRD-1, the *C. elegans* ortholog of STRAD, display impaired SIS

We found *strd-1* mutants consistently show impaired SIS. The gene *strd-1* encodes the *C. elegans* ortholog of STE20 related adaptor protein (STRAD), a conserved pseudokinase that functions with the kinase PAR-4/LKB1 in stress-responsive pathways (Narbonne et al., 2010). We tested two independent deletion, likely-null, alleles, *strd-1(ok2275)* and *strd-1(ok2283)* (Fig. 3C and D). Prior to UV exposure, *strd-1(ok2275)* showed higher median quiescence than N2, while *strd-1(ok2283)* shows lower median quiescence than N2. After the UV exposure, *strd-1(ok2275)* and *strd-1(ok2283)* show 34% and 12% reductions, respectively, in the median quiescence fraction compared with N2.

### Mutants for the active zone protein CLA-1 display elevated SIS

The gene c*la-1* encodes Clarinet, an active zone protein required for synaptic vesicle clustering and release (Krout et al., 2023; Xuan et al., 2017). We examined five independent mutant alleles (Fig. 3E and F), including *cla-1(gk352)*, *cla-1(ok2285)*, *cla-1(ok560)*, *cla-1(ok618)*, and *cla-1(ok937)*. Four of these five alleles contain deletions in the *cla-1* gene, suggesting that they are null alleles of *cla-1*. All the alleles except for *gk352* showed spontaneous quiescence not significantly higher than N2. After UV exposure, all 5 mutant alleles showed a higher median quiescence fraction than N2. On average, *cla-1* mutants show a 12% increase in post-UV quiescence compared with N2.

### Mutants for EGL-8, the *C. elegans* homolog of phospholipase C beta (PLCβ), show elevated SIS

The *egl-8* gene encodes the *C. elegans* homolog of phospholipase C beta (PLCβ), which is expressed throughout the nervous system as well as near intestinal cell junctions (Miller et al., 1999). We tested five independent alleles for *egl-8* (Fig. 3G and H), including *egl-8(e2917)*, *egl-8(md1971)*, *egl-8(n488)*, *egl-8(nj77)*, and *egl-8(sa47)*. Before the UV exposure, all *egl-8* alleles except *e2917* show quiescence not significantly higher than N2. After UV exposure, we observed an elevated median quiescence fraction in all the mutants compared with N2. All the alleles show at least a 38% increase in post-UV quiescence compared with N2.

## Discussion

In this work, we have demonstrated a semi-automated genetic screen for regulators of SIS in *C. elegans* using the MMP strain collection (Thompson et al., 2013). There are four main novel aspects to our study: (1) Robotic-assisted automation to increase throughput for genetic discovery; (2) Our focus on genes with moderate effects on sleep; and (3) Identifying genes whose disruption causes either a decrease or an increase in sleep. (4) The insight gained by identifying three novel genes regulating *C. elegans* sleep.

To improve the screening throughput, we performed automated imaging on the WorMotel substrates and applied robotic automation for animal manipulation. Some practical constraints, however, limited the proportion of the strains that could be screened using the robotic workflow (Results and Table S1). The screen was conducted jointly by two laboratories, and the robotic system was available in only one laboratory. In addition, the screen was initially performed manually and was later adapted to the robotic workflow. These constraints resulted in a larger portion of the strains being screened manually (Results and Table S1). Nevertheless, the robotic automation increased the screening throughput by ∼50% compared to the pure manual method (Materials and methods). In addition to increasing speed, the robotic system has the advantage of being able to work continuously without fatigue and reduces the variability of experimental operations. Our screening pipeline serves as a model for accelerating other genetic screens through automation technologies.

The computational method we used here, SKAT (Timbers et al., 2016), has important advantages over the classical genetic mapping methods. To maximize the chance of successful mapping, classical methods typically pick strains showing extreme phenotypes and perform genetic mapping on those strains. Examples of such large effect size mutants include *aptf*-1 (Turek et al., 2013), *ceh-17* (Hill et al., 2014), *goa-1* (Huang et al., 2017b), and *dgk-1* (Chen et al., 2023). Mutants causing moderate changes in the SIS phenotype are usually not prioritized for genetic mapping. However, analysis of mutants with moderate effects may be more relevant to clinical studies of sleep disorders, which are more likely to show a quantitative defect in sleep than to show a complete lack of sleep. Indeed, studies of naturally short sleepers generally show quantitative reductions but never complete elimination in sleep (Chen et al., 2025; He et al., 2009; Shi et al., 2019; Shi et al., 2021; Xing et al., 2019).

SKAT is capable of simultaneously taking into account the phenotypic and genomic data from a large number of strains, making it a powerful method for detecting mutants with moderate effect sizes, for example *cla-1* and *strd-1* identified in this work. Notably, the SKAT analysis can find applications in other studies using sequenced strain libraries (Crombie et al., 2024; Thompson et al., 2013), for example, in genetic screens related to neuronal function (Timbers et al., 2016), mechanosensory pathways (Lawry, 2020), and compound screen for anthelmintics (Mathew et al., 2016).

The candidate genes highlighted by the SKAT include three validated novel genetic regulators of SIS in *C. elegans*: *strd-1*, *cla-1*, and *egl-8*. While mutations in *strd-1* result in reduced SIS, mutations in *cla-1* and for *egl-8* are associated with an increase in SIS. With rare exceptions (Sinner et al., 2021), prior forward genetic studies in *C. elegans* have focused on mutations associated with reduced quiescence. Expanding our focus to include genes whose disruption causes increased quiescence, holds promise to deepen our understanding of sleep regulation. While genes whose disruption causes reduced quiescence may be relevant to clinical insomnia, genes whose disruption causes increased quiescence may be relevant to clinical hypersomnia.

A prior *C. elegans* study showed that STRD-1/STRAD promotes PAR-4/LKB1-dependent phosphorylation of AAK-2/AMPK (adenosine monophosphate-activated protein kinase) under metabolic stress (Narbonne et al., 2010). Activated AMPK inhibits anabolic pathways and activates catabolic pathways (Hardie, 2011), thereby shifting cellular metabolism toward energy production. The salt-induced kinase KIN-29/SIK acts in sensory neurons to promote SIS by activating the sleep-promoting ALA and RIS neurons (Grubbs et al., 2020). Together, these findings support a model in which STRD-1/STRAD functions with PAR-4/LKB1 to activate AAK-2/AMPK, and this metabolic signal upregulates the KIN-29/SIK pathway in sensory neurons upstream of ALA and RIS (Fig. 4A). According to this model, loss of STRD-1 would reduce AAK-2/AMPK activation, weaken KIN-29/SIK-dependent sleep signaling, and suppress SIS.

**Fig. 4.**
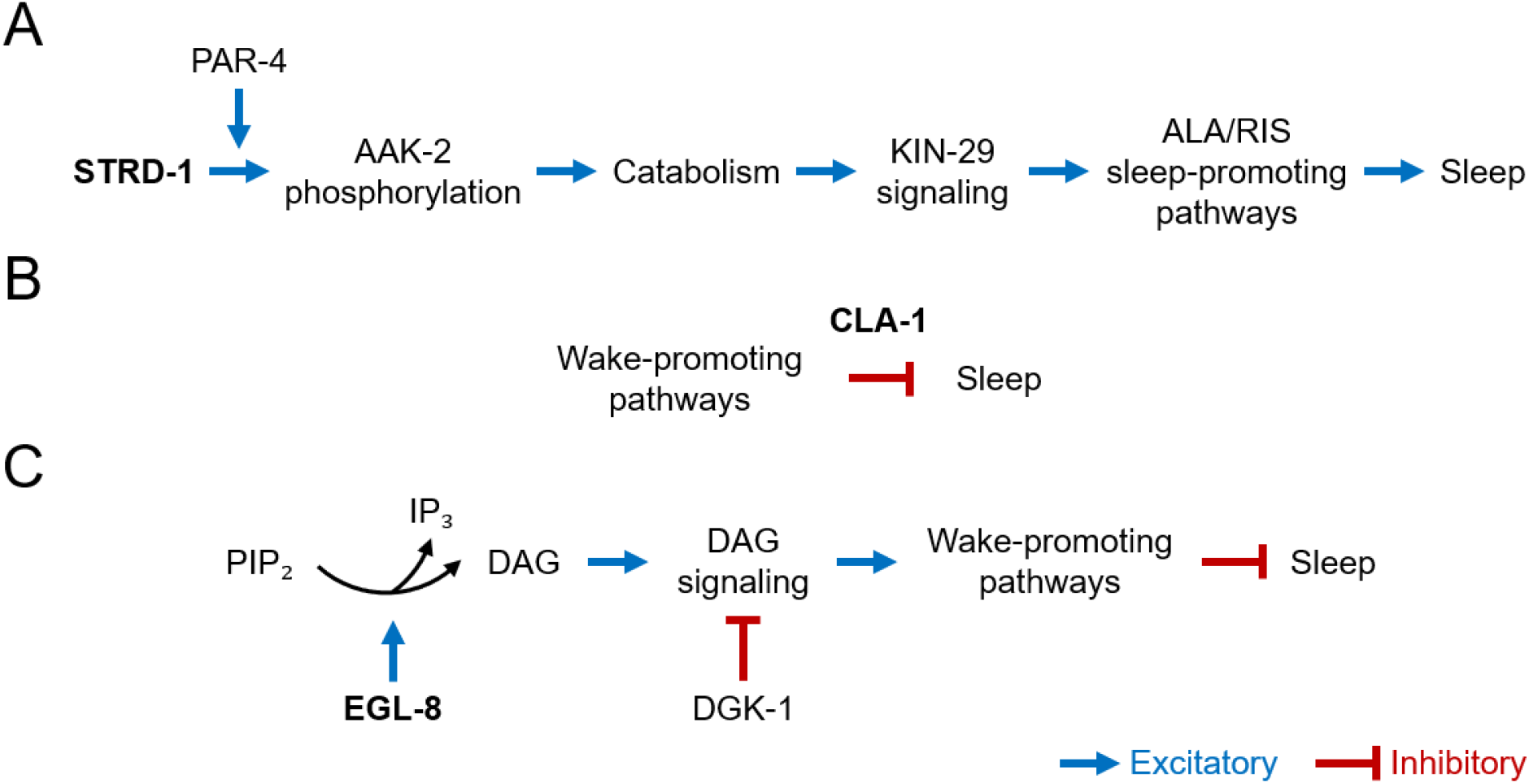
Speculative models of SIS regulation by the genes identified in this study. Potential models for roles of (A) STRD-1, (B) CLA-1, and (C) EGL-8 in SIS regulation.

CLA-1/Clarinet is a presynaptic active-zone protein required for synaptic vesicle clustering, active-zone organization, and sustained synaptic release (Krout et al., 2023; Xuan et al., 2017). Loss of CLA-1 disrupts vesicle clustering and increases synaptic depression (Krout et al., 2023; Xuan et al., 2017). CLA-1 regulates presynaptic autophagy (Xuan et al., 2023), suggesting a role in maintaining synaptic homeostasis. Based on these findings, we speculate that increased SIS in *cla-1* mutants (Fig. 3E and F) may reflect impaired function of arousal-or recovery-promoting neural circuits after stress (Fig. 4B). Mutant CLA-1 could weaken sustained neurotransmitter release, thereby reducing the ability of sensory neurons, interneurons, or motor circuits to emerge from quiescence after stress. In parallel, defective CLA-1-dependent presynaptic autophagy may increase synaptic stress or delay synaptic recovery following stress, further biasing animals toward a prolonged quiescent state.

EGL-8 is a phospholipase C beta (PLCβ) that functions downstream of the G protein EGL-30/Gqα and is broadly expressed in the nervous system (Miller et al., 1999). As a PLCβ, EGL-8 cleaves phosphatidylinositol 4,5-bisphosphate (PIP₂) into inositol 1,4,5-trisphosphate (IP₃) and diacylglycerol (DAG) (Jia et al., 2024; Miller et al., 1999). A prior study found that increased DAG signaling can suppress sleep in *C. elegans* (Chen et al., 2023). Specifically, for DAG kinase mutants *dgk-1*, in which the DAG signaling is insufficiently restrained, *C. elegans* shows reduction in both DTS and SIS (Chen et al., 2023). Together, these findings suggest a model in which EGL-8 regulates SIS by modulating DAG-dependent signaling, which is normally negatively regulated by DGK-1 (Fig. 4C). Loss of EGL-8 function may reduce DAG signaling and thereby promote increased SIS.

### Limitations of the study

In this study, hits were validated based on consistent SIS phenotypes observed across 2 or more independent mutant alleles. Additional experiments, such as complementation testing, genetic rescue, genetic linkage mapping, and CRISPR-based genome editing, would further support causal roles of these genes in influencing the SIS phenotype.

Our current SKAT analysis for estimating the associations between the genes and sleep phenotypes treat all types of non-synonymous mutations equally. Previous suggests that assigning weights to different types of mutations could increase the power of the analysis (Timbers et al., 2016). For example, computational tools such as SIFT (Kumar et al., 2009) and Polyphen (Adzhubei et al., 2010) could be used to estimate the severity of each mutation.

Although this work identifies new genes associated with changes in SIS phenotypes, we did not provide mechanistic evidence for how these genes are involved in SIS regulation. Nevertheless, the speculative models we proposed here may guide future studies to characterize their roles (Fig. 4).

## Supporting information

Software S1

Tabls S2

## Data availability

Information on the strains used in this study, including their sleep scores and availability, can be found in Table S1. The ranks of individual known sleep-regulating genes from the SKAT analysis are available in Table S2. The full lists of ranked genes from the SKAT analysis with *N_a_* = 5 and *N_a_* = 3 can be found in Tables S3 and S4, respectively. The data files and source code used to reproduce the figures and SKAT analysis are available in Software S1.

## Acknowledgements

Some of the *C. elegans* strains were provided by the *Caenorhabditis* Genetics Center, which is funded by the NIH Office of Research Infrastructure Programs (P40 OD010440). Some *C. elegans* strains were provided by the National BioResource Project of Japan. We thank Helen Chamberlin, Erik Andersen, Animesh Biswas, and Alex Ng Tung Hing for helpful discussions and suggestions.

