## Supplementary material for "A semi-automated genetic screen using genome-wide association analysis identifies modulators of stress-induced sleep in *Caenorhabditis elegans*": Software S1: Main_figure1.m

```
%% Script for plotting Figure 1 of the Sleep Screen paper.
% Panels: Fig1B (activity heatmap), Fig1C (quiescence-fraction heatmap),
%         Fig1D (quiescence-fraction scatter for N2 / aptf-1 / ceh-17).
%
% Helper functions (fracQ, fracQbin230817, beeswarm) live in ../helpers/.

clear all;
close all;
clc;

addpath(fullfile('..', 'helpers'));

load('data.mat');

%% ratio for pbaspect ratio
ratio = 1;

%% === Fig1B ===
close all;

gpLab = strings(1,141);                 % 1×48 of ""
gpLab([24 72 122]) = ["N2","\it aptf-1(tm3287)","\it ceh-17(np1)"];  % set only those ticks

figure
qPRE = datPRE';
qPOST = datPOST';
qALL=[qPRE(1:10:end-1,:); qPOST(1:10:end,:)];
imagesc(qALL') %make heatmap showing fracQ data
title('Activity')
xticks(1:36:size(qALL,1))         % ticks every hour
xticklabels(0:4)
xlabel('Time (hour)')
yticks([])
ax = gca;
ax.FontSize = 30;

colorbar
colormap hot

% === draw 3 white lines every 16 rows ===
hold on
ax.Layer = 'top';                  % keep lines on top of image
nRows = size(qALL,2);              % = 48 (rows after transpose)
blk   = 48;
edges = blk:blk:nRows-1;             % [16 32 48]
for k = edges
    y = k + 0.5;                   % boundary between row k and k+1
    if y <= nRows + 0.5
        plot(xlim, [y y], 'w-', 'LineWidth', 5);
    end
end

hold off

pbaspect([ratio 1 1]);

f = gcf;
f.Units = 'pixels';
f.Position = [100 100 800 800/ratio];

exportgraphics(gca, "Fig1B.svg");

%% === Fig1C ===
close all;
figure
[qPRE,t3PRE,p3PRE]=fracQ(datPRE',1,10,60);
[qPOST,t3POST,p3POST]=fracQ(datPOST',1,10,60);
qALL=[qPRE(1:10:end,:); qPOST(1:10:end,:)];

imagesc(qALL')                     % heatmap
title('Quiescence fraction')
xticks(1:360:size(qALL,1))         % ticks every hour
xticklabels(0:4)
xlabel('Time (hour)')
yticks([])
ax = gca; ax.FontSize = 30;
colorbar; colormap jet

% === draw 3 white lines every 16 rows ===
hold on
ax.Layer = 'top';                  % keep lines on top of image
nRows = size(qALL,2);              % = 48 (rows after transpose)
blk   = 48;
edges = blk:blk:nRows-1;             % [16 32 48]
for k = edges
    y = k + 0.5;                   % boundary between row k and k+1
    if y <= nRows + 0.5
        plot(xlim, [y y], 'w-', 'LineWidth', 5);
    end
end
hold off

pbaspect([ratio 1 1]);

f = gcf;
f.Units = 'pixels';
f.Position = [100 100 800 800/ratio];

exportgraphics(gca, "Fig1C.svg");

%% === Fig1D ===
close all;

% Scatterplot of total quiescence fraction

%ENTER wells to be censored
cens = [14, 17, 32, 34, 48, 50, 53, 61, 71, 108, 109, 131, 144]; % N2, N2, N2, aptf-1, aptf-1, aptf-1, ceh-17, ceh-17, ceh-17

%ENTER THE HOW MANY STRAINS
N_STRAINS = 3;

%condition names will be entered based on strains provided above
%(can be copy/pasted directly to prism
gpNames=cell(N_STRAINS,1);
gpNames{1}=["N2"];
gpNames{2}=["\it aptf-1(tm3287)"];
gpNames{3}=["\it ceh-17(np1)"];

%ENTER wells for corresponding conditions
%list which animals go in which group

% a vector store the begin/end well number for each group
% the length of the vector must be an even number, since there are two
% number for each group, begin well number and end well number.
% the pattern: [begin gp 1, end gp 1; begin gp 2, end gp 2; ...]
well_num_begin_end = [1,48;49,96;97,144];

gp = cell(1, N_STRAINS);
gp{1}=[well_num_begin_end(1,1) : well_num_begin_end(1,2)]; %STR A
gp{2}=[well_num_begin_end(2,1) : well_num_begin_end(2,2)]; %STR B
gp{3}=[well_num_begin_end(3,1) : well_num_begin_end(3,2)]; %STR C
gpN=length(gp); %calculate number of groups
gpLens=zeros(1,gpN); %N per group (filled below)
nOrig=zeros(1,gpN); %number of animals in original groups
nCens=zeros(1,gpN); %number of animals after removing censored animals
for i = 1:length(gp)
    nOrig(i)=length(gp{i});

    gp{i}=setdiff(gp{i},cens); %remove censored wells from groups
    gpLens(i)=length(gp{i}); %get number of animals in each group

    nCens(i)=nOrig(i)-gpLens(i);
end
clear i
gpMax=max(gpLens); %find largest group
TimepImg = 10;
EPM= 60 / TimepImg; %epochs per min (1 image/10 seconds)
groupSz=48; %size of groups for input into prism (standard size makes it easier to copy onto preexisting files)

%out3 qTotal - sum of quiescence over all 4 hours
%use prism grouped input for 2-way anova design
%so that genotype and treatment are matched appropriately across groups

%same set up as out1, except sum over 4 hours

ATp=0.01; %activity threshold: fraction of 95th percentile of activity
QT=0.25; %qui threshold: fraction of moving window needed to be quiescent
qWin=60; %qWin is moving window size (in number of frames)
binTime = 60; % binTime time duration (min) of each bin window
binSz = binTime * 4 * EPM; %binSize here refers to number of frames in each bin

[qTime,~,~,qTimePRE,~,durPre] = fracQbin230817(datPRE',datPOST',gp,ATp,QT,qWin,binSz,EPM);

nCol=length(gp); %number of groups of columns in output

out3_qTot=NaN(2,groupSz*nCol);
for iCols = 1:nCol
    colSt=(iCols-1) * groupSz + 1;


    quiPRE=qTimePRE{iCols} * (EPM/binSz);
    quiPOST=qTime{iCols} * (EPM/binSz);
    colEn=colSt+gpLens(iCols)-1;

    out3_qTot(1,colSt:colEn)=quiPRE;
    out3_qTot(2,colSt:colEn)=quiPOST;

end

figure

% flatten the data for beeswarm plot

% category vector w/o merge
cate_vec = [];

% pre-UV
QfTot_vec_preUV = [];

% post-UV
QfTot_vec_postUV = [];


% all
cate_vec_all = [];
QfTot_vec_all = [];

% color for plotting different groups
cmap = [];
cOrder = colororder;
Color_preUV = [0 0 0];
Color_postUV = cOrder(4,:);

for i = 1 : size(gp, 2)

    % data frame
    gp_QfTot = out3_qTot(:, well_num_begin_end(i, 1): well_num_begin_end(i, 2));

    % remove NaN
    gp_QfTot_preUV = rmmissing(gp_QfTot(1,:));
    gp_QfTot_postUV = rmmissing(gp_QfTot(2,:));

    % category vector
    cate_vec = [cate_vec, i * ones(1, size(gp_QfTot_postUV, 2))];

    % pre-UV
    QfTot_vec_preUV = [QfTot_vec_preUV, gp_QfTot_preUV];


    % add pre-UV to merged vector
    cate_vec_all = [cate_vec_all, (2 * i - 1) * ones(1, size(gp_QfTot_preUV, 2))];
    QfTot_vec_all = [QfTot_vec_all, gp_QfTot_preUV];

    cmap = [cmap; Color_preUV]; % plot color for pre-UV


    % post-UV
    QfTot_vec_postUV = [QfTot_vec_postUV, gp_QfTot_postUV];

    % add post-UV to merged vector
    QfTot_vec_all = [QfTot_vec_all, gp_QfTot_postUV];
    cate_vec_all = [cate_vec_all, (2 * i) * ones(1, size(gp_QfTot_postUV, 2))];

    cmap = [cmap; Color_postUV]; % use a different plot color with pre-UV

end

dot_sz = 3;

% odd categories = pre-UV (hollow triangle), even categories = post-UV (filled circle)
mkrs = repmat({'^','o'}, 1, size(gp,2));
mkrFill = repmat({false, true}, 1, size(gp,2));

xbee = beeswarm(cate_vec_all',QfTot_vec_all', ...
    'dot_size', dot_sz, ...
    'overlay_style','sd',...
    'corral_style','random',...
    'colormap',cmap, ...
    'Marker', mkrs, ...
    'MarkerFill', mkrFill, ...
    'MarkerLineWidth', 1.5, ...
    'MarkerFaceAlpha', 0.5, ...
    'MarkerEdgeColor', 'w');


N2_pre_UV = QfTot_vec_all(cate_vec_all == 1);
N2_post_UV = QfTot_vec_all(cate_vec_all == 2);
aptf_1_pre_UV = QfTot_vec_all(cate_vec_all == 3);
aptf_1_post_UV = QfTot_vec_all(cate_vec_all == 4);
ceh_17_pre_UV = QfTot_vec_all(cate_vec_all == 5);
ceh_17_post_UV = QfTot_vec_all(cate_vec_all == 6);

fprintf('N2 vs aptf-1\n')
ranksum(N2_pre_UV, aptf_1_pre_UV, 'tail', 'right')
ranksum(N2_post_UV, aptf_1_post_UV, 'tail', 'right')

fprintf('N2 vs ceh-17\n')
ranksum(N2_pre_UV, ceh_17_pre_UV, 'tail', 'right')
ranksum(N2_post_UV, ceh_17_post_UV, 'tail', 'right')

ylim([0, 1]);
xlim([0, 2 * size(gp, 2)+0.8]);

ylabel('Quiescence fraction')

xticks(1.5 : 2 : 2 * size(gp, 2));
xticklabels(gpNames)

y_leg_preUV = 0.90;
y_leg_postUV = 0.83;

% draw the legend
hold on
hold off

ax = gca; ax.FontSize = 33;
ax.LineWidth = 2;
xtickangle(20)

pbaspect([ratio 1 1]);

f = gcf;
f.Units = 'pixels';
f.Position = [100 100 800 800/ratio];

exportgraphics(gca, "Fig1D.svg");
```
