## Supplementary material for "A semi-automated genetic screen using genome-wide association analysis identifies modulators of stress-induced sleep in *Caenorhabditis elegans*": Software S1: Main_figure2.m

```
%% Script for plotting Figure 2 of the Sleep Screen paper.
% Panels: Fig2A (mean-quiescence histogram for MMP strains vs N2),
%         Fig2B (pre-UV vs post-UV scatter for MMPs and N2).
%
% Helper functions live in ../helpers/.

clear all;
close all;
clc;

addpath(fullfile('..', 'helpers'));

load('robot_MMP_Qf.mat')
load('sleep_data_cleaned.mat')

%% correct known strain-name errors in Kerry's dataset
strain_names(strcmpi(strain_names, 'VC20497C')) = {'VC20497'};
idx_VC20750N = strcmpi(strain_names, 'VC20750 N'); % duplicate mislabel of VC20750, drop it
strain_names(idx_VC20750N) = [];
stressed_quiescence_mean(idx_VC20750N) = [];
unstressed_quiescence_mean(idx_VC20750N) = [];

stressed_quiescence_mean_kerry = stressed_quiescence_mean'/60;
unstressed_quiescence_mean_kerry = unstressed_quiescence_mean'/60;
load('sleep_data_N2_Kerry')
load('robot_N2_Qf_byWM.mat')

%% remove duplicate strains in Kerry's dataset

[unique_MMP_Names_kerry, ~, idx] = unique(lower(strain_names'), 'stable');
unique_stressed_quiescence_mean_kerry = accumarray(idx(:), stressed_quiescence_mean_kerry(:), [], @mean);
unique_unstressed_quiescence_mean_kerry = accumarray(idx(:), unstressed_quiescence_mean_kerry(:), [], @mean);

%% colors
cOrder = colororder;
Color_postUV = cOrder(4,:);

%% === Fig2A : Plot the histogram of the mean quiescence for MMPs ===

close all;

% calculate the pre and post-UV for N2
N_N2_kerry_preUV = size(unstressed_quiescence_mean_N2_kerry, 2) * 8;
N_N2_kerry_postUV = size(stressed_quiescence_mean_N2_kerry, 2) * 8;

N_N2_john_preUV = size(Qf_all_ctrl_preUV, 1);
N_N2_john_postUV = size(Qf_all_ctrl, 1);

% Calculate the mean for pool N2s
meanQf_N2_preUV_pool = (N_N2_kerry_preUV * mean(unstressed_quiescence_mean_N2_kerry) + ...
    N_N2_john_preUV * mean(Qf_all_ctrl_preUV))/(N_N2_kerry_preUV + N_N2_john_preUV);

meanQf_N2_postUV_pool = (N_N2_kerry_postUV * mean(stressed_quiescence_mean_N2_kerry) + ...
    N_N2_john_postUV * mean(Qf_all_ctrl))/(N_N2_kerry_postUV + N_N2_john_postUV);


merge_mean_Qf_preUV = [unique_unstressed_quiescence_mean_kerry;ranked_mean_Qf_preUV];
merge_mean_Qf_postUV = [unique_stressed_quiescence_mean_kerry;ranked_mean_Qf];


% remove strain having both 0 pre and post UV
idx_to_remove = (merge_mean_Qf_preUV == 0) .* (merge_mean_Qf_postUV == 0);

merge_mean_Qf_preUV_filtered = merge_mean_Qf_preUV(~idx_to_remove);
merge_mean_Qf_postUV_filtered = merge_mean_Qf_postUV(~idx_to_remove);

%% === Supplemental table : per-strain pre/post-UV quiescence underlying Fig2A/B ===

ranked_MMP_name_list_clean = erase(cellstr(ranked_MMP_name_list), ' UV');
merge_MMP_Names = upper([unique_MMP_Names_kerry; ranked_MMP_name_list_clean]);
merge_MMP_Names_filtered = merge_MMP_Names(~idx_to_remove);

% unique_MMP_Names_kerry strains were screened by a human; the rest by the robot
merge_Screen_Method = [repmat({'human'}, numel(unique_MMP_Names_kerry), 1); ...
    repmat({'robot'}, numel(ranked_MMP_name_list_clean), 1)];
merge_Screen_Method_filtered = merge_Screen_Method(~idx_to_remove);

Genotype = repmat({'MMP'}, numel(merge_MMP_Names_filtered), 1);
Identifier = strcat('https://cgc.umn.edu/strain/', merge_MMP_Names_filtered);
Source = repmat({'Caenorhabditis Genetics Center'}, numel(merge_MMP_Names_filtered), 1);

supp_table_Fig2 = table(merge_MMP_Names_filtered, Genotype, Identifier, Source, ...
    merge_Screen_Method_filtered, merge_mean_Qf_preUV_filtered, merge_mean_Qf_postUV_filtered, ...
    'VariableNames', {'Strain_Name', 'Genotype', 'Identifier', 'Source', ...
    'Screened_by', 'PreUV_Quiescence_Fraction', 'PostUV_Quiescence_Fraction'});
writetable(supp_table_Fig2, 'Fig2_MMP_quiescence_data.csv');

% Histogram of the mean quiescence for MMPs
edges= [0:0.03:1];
figure();
hold on;

N2_postUV_Qf_all = [stressed_quiescence_mean_N2_kerry'; robot_N2_Qf_postUV_byWM'];
histogram(N2_postUV_Qf_all, ...
    edges, 'Normalization', 'probability', ...
    'FaceColor', 'g', 'FaceAlpha', 0.4, 'EdgeColor', 'g', 'EdgeAlpha', 1, ...
    'LineStyle','none')


N2_preUV_Qf_all = [unstressed_quiescence_mean_N2_kerry'; robot_N2_Qf_preUV_byWM'];
histogram(N2_preUV_Qf_all, ...
    edges, 'Normalization', 'probability', ...
    'FaceColor', [0.8 0.2 0], 'FaceAlpha', 0.4, 'EdgeColor', [0.8 0.2 0], 'EdgeAlpha', 1, ...
    'LineStyle','none')


histogram(merge_mean_Qf_preUV_filtered, edges, 'Normalization', 'probability', ...
    'FaceColor', 'k', 'FaceAlpha', 0, 'EdgeColor', 'k', 'EdgeAlpha', 1,  ...
    'LineStyle','--', 'LineWidth', 3)
histogram(merge_mean_Qf_postUV_filtered, edges, 'Normalization', 'probability',...
    'FaceColor', Color_postUV, 'FaceAlpha', 0, 'EdgeColor', Color_postUV, 'EdgeAlpha', 1, ...
    'LineStyle','-', 'LineWidth', 3)


hold off;


xlabel('Mean quiescence fraction')
ylabel('Relative frequency')

ax = gca; ax.FontSize = 25;
ax.LineWidth = 2;
pbaspect([1 1 1]);

xticks(0 : 0.2 : 1);

xlim([0 1])

f = gcf;
f.Units = 'pixels';
f.Position = [100 100 800 800];
box off;

exportgraphics(gca, "Fig2A.svg");

% test of variance
disp('test of variance for N2 and MMP')
var_test_data_postUV = [merge_mean_Qf_postUV_filtered; N2_postUV_Qf_all];
groups_data_postUV = [ones(size(merge_mean_Qf_postUV_filtered)); 2 * ones(size(N2_postUV_Qf_all))];
vartestn(var_test_data_postUV, groups_data_postUV, 'TestType', 'BrownForsythe');

var_test_data_preUV = [merge_mean_Qf_preUV_filtered; N2_preUV_Qf_all];
groups_data_preUV = [ones(size(merge_mean_Qf_preUV_filtered)); 2 * ones(size(N2_preUV_Qf_all))];
vartestn(var_test_data_preUV, groups_data_preUV, 'TestType', 'BrownForsythe');

% test of mean
disp('test of mean for N2 and MMP')
p = ranksum(merge_mean_Qf_postUV_filtered,N2_postUV_Qf_all, 'tail', 'right')
p = ranksum(merge_mean_Qf_preUV_filtered,N2_preUV_Qf_all, 'tail', 'right')

%% === Fig2B : Scatter plot of the mean quiescence for MMPs ===
figure();
hold on;
scatter(merge_mean_Qf_preUV_filtered, merge_mean_Qf_postUV_filtered, ...
    'filled', 'MarkerFaceColor', 0.5*Color_postUV + 0.5*[0 0 0], 'MarkerFaceAlpha', 0.6);
scatter(N2_preUV_Qf_all, N2_postUV_Qf_all, 'filled', 'MarkerFaceColor', 'g', 'MarkerFaceAlpha', 0.2);

xlabel('Mean pre-UV quiescence fraction')
ylabel('Mean post-UV quiescence fraction')
ax = gca; ax.FontSize = 25;
ax.LineWidth = 2;
xlim([0 1])
ylim([0 1])

xl = xlim;
plot(xl, xl, '--k', 'LineWidth', 2)

f = gcf;
f.Units = 'pixels';
f.Position = [100 100 800 800];
pbaspect([1 1 1]);

hold off
exportgraphics(gca, "Fig2B.svg");
```
