## Supplementary material for "A semi-automated genetic screen using genome-wide association analysis identifies modulators of stress-induced sleep in *Caenorhabditis elegans*": Software S1: Main_figure3.m

```
%% Script for plotting Figure 3 of the Sleep Screen paper.
% Panels:
%   Fig3A : MMP-vs-N2 histogram (per-wormotel N2 reconstruction).
%   Fig3B : Pre-UV vs post-UV mean ± SEM dot plot for follow-up alleles.
%   Fig3C-H (per-gene) : trace heatmap + beeswarm scatter for strd-1, cla-1, egl-8
%                        (C/D = strd-1, E/F = cla-1, G/H = egl-8; heatmap/scatter).
%
% Helper functions (fracQ, beeswarm) live in ../helpers/.
%
% The per-gene block loads figure3/data_per_gene.mat, a lightweight
% preprocessed file. To regenerate it from raw recordings, edit and run
% preprocess_per_gene_data.m (paths there must point at your local copy
% of the raw data).

clear all;
close all;
clc;

addpath(fullfile('..', 'helpers'));

%% === Fig3A : MMP-vs-N2 histogram ===
clear; clc; close all;

% Only resultsTable/strainInfo are used below; data_strain_summary.mat is
% trimmed to just these two variables to keep the file small.
load('data_strain_summary.mat', 'resultsTable', 'strainInfo');

%% === SKAT percentile lookup maps ===
% SKAT_filtered_Na5.tsv is the SKAT gene-level association table (see
% ../SKAT/README.md), filtered to N.Marker.Test >= 5, ranked and
% converted to a percentile. Lives alongside this script.
skat = readtable('SKAT_filtered_Na5.tsv', 'FileType', 'text', 'Delimiter', '\t');
skatSetID = string(skat.SetID);
skatCommon = string(skat.Common_name);

byCommon = containers.Map('KeyType', 'char', 'ValueType', 'double');
bySetid  = containers.Map('KeyType', 'char', 'ValueType', 'double');

% Canonical-gene-ID maps (SetID is unique per gene; Common_name is not
% always present), used to count unique genes regardless of whether a
% strain's Genotype references the gene by Common_name or SetID.
commonToSetid = containers.Map('KeyType', 'char', 'ValueType', 'char');
setidToSetid  = containers.Map('KeyType', 'char', 'ValueType', 'char');

for i = 1:height(skat)
    bySetid(char(skatSetID(i))) = skat.Percentile(i);
    setidToSetid(char(skatSetID(i))) = char(skatSetID(i));
    if skatCommon(i) ~= "N.A." && ~isKey(byCommon, char(skatCommon(i)))
        byCommon(char(skatCommon(i))) = skat.Percentile(i);
        commonToSetid(char(skatCommon(i))) = char(skatSetID(i));
    end
end

% Genotype gene names that don't match the tsv's SetID/Common_name
% directly (confirmed by manual lookup on WormBase), mapped to the
% identifier actually used in the tsv.
geneAliases = containers.Map( ...
    {'F32E10.7', 'crmb-1'}, ...
    {'cla-1', 'T21D12.11'});

%% Filtered table only for strain-level histogram
% Convert to string first (safe for manipulation)
strainInfo.StrainName = string(strainInfo.StrainName);

% Add " UV" to each strain
strainInfo.StrainName = strainInfo.StrainName + " UV";

% Remove any accidental spaces
strainInfo.StrainName = strtrim(strainInfo.StrainName);

% Convert to categorical
strainInfo.StrainName = categorical(strainInfo.StrainName);


strainInfo.Genotype = categorical(cellstr(strainInfo.Genotype));

% Drop strains not from the SKAT priority list, except N2 (always kept as
% the control). This removes them from the histogram, the
% variance/rank-sum tests below, and the CSV in one place.
hasSkatMatch = false(height(strainInfo), 1);
for i = 1:height(strainInfo)
    pct = lookupSkatPercentile(char(string(strainInfo.Genotype(i))), byCommon, bySetid, geneAliases);
    hasSkatMatch(i) = ~isnan(pct);
end
isN2Strain = strainInfo.StrainName == "N2 UV";
strainInfo = strainInfo(hasSkatMatch | isN2Strain, :);

resultsTable2 = resultsTable(ismember(resultsTable.Strain, strainInfo.StrainName), :);

%% Mean per strain (for histogram)
[G, groupNames] = findgroups(resultsTable2.Strain);
mean_pre  = splitapply(@mean, resultsTable2.Qf_pre, G);
mean_post = splitapply(@mean, resultsTable2.Qf_post, G);

%% === Supplemental table : per-strain pre/post-UV quiescence underlying Fig3A ===
strain_names_fig3 = erase(cellstr(groupNames), ' UV');

% Genotype/Identifier/Source looked up from a local copy of Table_S1's
% candidate_screen tab (candidate_screen_strain_info.csv), so this script
% does not depend on a file outside the repo.
candidateScreen = readtable('candidate_screen_strain_info.csv');

[isFound, locS1] = ismember(lower(strain_names_fig3), lower(candidateScreen.StrainName));
Genotype = repmat({''}, numel(strain_names_fig3), 1);
Identifier = repmat({''}, numel(strain_names_fig3), 1);
Source = repmat({''}, numel(strain_names_fig3), 1);
Genotype(isFound) = candidateScreen.Genotype(locS1(isFound));
Identifier(isFound) = candidateScreen.Identifier(locS1(isFound));
Source(isFound) = candidateScreen.Source(locS1(isFound));

% One percentile per matched gene locus, semicolon-joined for multi-locus
% genotypes (e.g. double mutants); loci with no SKAT match are simply
% omitted rather than shown as "NA". "NA" is only used when no locus in
% the genotype matches at all (e.g. N2, which has no mutant gene).
SKAT_Rank_Percentile = strings(numel(Genotype), 1);
for i = 1:numel(Genotype)
    pctList = lookupSkatPercentiles(Genotype{i}, byCommon, bySetid, geneAliases);
    matchedPct = pctList(~isnan(pctList));
    if isempty(matchedPct)
        SKAT_Rank_Percentile(i) = "NA";
    else
        SKAT_Rank_Percentile(i) = strjoin(string(matchedPct), "; ");
    end
end

supp_table_Fig3 = table(strain_names_fig3, Genotype, Identifier, Source, mean_pre, mean_post, SKAT_Rank_Percentile, ...
    'VariableNames', {'Strain_Name', 'Genotype', 'Identifier', 'Source', ...
    'PreUV_Quiescence_Fraction', 'PostUV_Quiescence_Fraction', 'SKAT_Rank_Percentile'});
writetable(supp_table_Fig3, 'Fig3_mutants_quiescence_data.csv');

%% === Unique gene count in the Fig3 CSV ===
% Canonicalize each strain's SKAT-matched gene(s) to a SetID and count
% distinct genes, so different alleles of the same gene (e.g. AR10/AR11/
% AR4, all cysl-1) collapse to one entry. N2 and any unmatched strain
% contribute no gene. A multi-locus genotype (e.g. VC2725) can contribute
% more than one gene.
allCanonicalGenes = strings(0,1);
strainOfGene = strings(0,1);
for i = 1:numel(Genotype)
    ids = resolveGeneID(Genotype{i}, commonToSetid, setidToSetid, geneAliases);
    allCanonicalGenes = [allCanonicalGenes; ids]; %#ok<AGROW>
    strainOfGene = [strainOfGene; repmat(string(strain_names_fig3{i}), numel(ids), 1)]; %#ok<AGROW>
end
[uniqueGenes, ~, geneGroupIdx] = unique(allCanonicalGenes);
fprintf('Fig3 CSV: %d strains represent %d unique gene(s)\n', numel(Genotype), numel(uniqueGenes));

fprintf('Genes represented by multiple strains:\n');
for k = 1:numel(uniqueGenes)
    strainsForGene = strainOfGene(geneGroupIdx == k);
    if numel(strainsForGene) > 1
        fprintf('  %-12s : %s\n', uniqueGenes(k), strjoin(strainsForGene, ', '));
    end
end

%% Reconstruct N2 UV wormotels from ORIGINAL unfiltered table
dates = unique(resultsTable.Date);

n2_mean_pre_by_wormotel  = [];
n2_mean_post_by_wormotel = [];
n2_block_sizes           = [];
n2_block_date            = strings(0,1);

for d = 1:numel(dates)

    thisDate = dates(d);
    Tdate = resultsTable(resultsTable.Date == thisDate, :);

    % Find N2 UV rows within this date in the ORIGINAL table
    n2_rows = find(Tdate.Strain == "N2 UV");

    if isempty(n2_rows)
        continue;
    end

    % Split into contiguous blocks
    breaks = [1; find(diff(n2_rows) > 1) + 1];
    numBlocks = numel(breaks);

    for b = 1:numBlocks

        startIdx = breaks(b);

        if b < numBlocks
            endIdx = breaks(b+1) - 1;
        else
            endIdx = numel(n2_rows);
        end

        blockRows = n2_rows(startIdx:endIdx);
        blockSize = numel(blockRows);

        n2_block_sizes(end+1,1) = blockSize;
        n2_block_date(end+1,1)  = string(thisDate);

        n2_mean_pre_by_wormotel(end+1,1) = ...
            mean(Tdate.Qf_pre(blockRows), 'omitnan');

        n2_mean_post_by_wormotel(end+1,1) = ...
            mean(Tdate.Qf_post(blockRows), 'omitnan');
    end
end

%% Plot
cOrder = colororder;
Color_postUV = cOrder(4,:);

figure;
hold on;
edges= [0:0.03:1];

% Histogram of N2 wormotel means (foreground)

histogram(n2_mean_pre_by_wormotel,...
    edges, 'Normalization', 'probability', ...
    'FaceColor', [0.8 0.2 0], 'FaceAlpha', 0.4, 'EdgeColor', [0.8 0.2 0], 'EdgeAlpha', 1, ...
    'LineStyle','none')

histogram(n2_mean_post_by_wormotel, ...
    edges, 'Normalization', 'probability', ...
    'FaceColor', 'g', 'FaceAlpha', 0.4, 'EdgeColor', 'g', 'EdgeAlpha', 1, ...
    'LineStyle','none')


% Histogram across strains (background)
histogram(mean_pre, edges, 'Normalization', 'probability', ...
    'FaceColor', 'k', 'FaceAlpha', 0, 'EdgeColor', 'k', 'EdgeAlpha', 1,  ...
    'LineStyle','--', 'LineWidth', 3);

histogram(mean_post, edges, 'Normalization', 'probability',...
    'FaceColor', Color_postUV, 'FaceAlpha', 0, 'EdgeColor', Color_postUV, 'EdgeAlpha', 1, ...
    'LineStyle','-', 'LineWidth', 3)

hold off;

xlabel('Mean quiescence fraction');
ylabel('Relative frequency');

ax = gca;
ax.FontSize = 25;
ax.LineWidth = 2;
pbaspect([1 1 1]);

xticks(0:0.2:1);
xlim([0 1]);

f = gcf;
f.Units = 'pixels';
f.Position = [100 100 800 800];

box off;

exportgraphics(gca, "Fig3A.svg");

%% Test of variance and mean

% test of variance
disp('test of variance for N2 and Mutants')
var_test_data_postUV = [mean_post; n2_mean_post_by_wormotel];
groups_data_postUV = [ones(size(mean_post)); 2 * ones(size(n2_mean_post_by_wormotel))];
vartestn(var_test_data_postUV, groups_data_postUV, 'TestType', 'BrownForsythe');

var_test_data_preUV = [mean_pre; n2_mean_pre_by_wormotel];
groups_data_preUV = [ones(size(mean_pre)); 2 * ones(size(n2_mean_pre_by_wormotel))];
vartestn(var_test_data_preUV, groups_data_preUV, 'TestType', 'BrownForsythe');

% test of mean
disp('test of mean for N2 and Mutants')
p = ranksum(mean_post, n2_mean_post_by_wormotel, 'tail', 'right')
p = ranksum(mean_pre, n2_mean_pre_by_wormotel, 'tail', 'right')

%% === Fig3B : Pre vs Post-UV per-strain dot plot ===
clear; clc; close all;

addpath(fullfile('..', 'helpers'));

load('data_per_worm.mat');

cOrder = colororder;
Color_postUV = cOrder(4,:);

% === Aggregate per strain ===
[G, strainList] = findgroups(Strain_names_all);

% --- Quiescence ---
preUV_Qf_mean = splitapply(@mean, preUV_Qf_all, G);
postUV_Qf_mean = splitapply(@mean, postUV_Qf_all, G);
SE_pre_Qf_mean = splitapply(@(x) std(x)/sqrt(numel(x)), preUV_Qf_all, G);
SE_post_Qf_mean = splitapply(@(x) std(x)/sqrt(numel(x)), postUV_Qf_all, G);

% === Identify N2 and reorder so it plots last ===
isN2 = contains(string(strainList), 'N2', 'IgnoreCase', true);
order = [find(~isN2); find(isN2)];

strainList = strainList(order);

% Quiescence
preUV_Qf_mean  = preUV_Qf_mean(order);
postUV_Qf_mean = postUV_Qf_mean(order);
SE_pre_Qf_mean  = SE_pre_Qf_mean(order);
SE_post_Qf_mean = SE_post_Qf_mean(order);


% === Define gene-group colors ===
geneColors = containers.Map( ...
    {'N2', 'strd-1', 'cla-1', 'egl-8','aptf-1', 'ceh-17'}, ...
    {[0.5 0.5 0.5], ...   % N2  (grey)
     [0.70, 0.00, 0.70], ...   % strd-1 (purple)
     [0.20, 0.90, 0.20], ...   % cla-1 (green)
     [1.00, 0.00, 0.00], ...   % egl-8 (red)
     [1.00, 0.50, 0.00], ...   % aptf-1 (orange)
     [0.15, 0.50, 0.85]} ...   % ceh-17 (blue)
);

% === Define shapes per gene ===
geneShapes = containers.Map( ...
    {'N2', 'strd-1','cla-1', 'egl-8', 'aptf-1', 'ceh-17'}, ...
    {'p', 's', '^', 'd', 'v', 'o'} ...
);

% === Strain → Gene mapping ===
strainToGene = containers.Map( ...
    {'N2 UV', ...
     'RB1771 UV', 'RB1775 UV', ...
     'VC387 UV', 'RB778 UV', 'RB1777 UV', 'VC596 UV', 'VC631 UV', ...
     'CB6614 UV', 'RM2221 UV', 'IK777 UV', 'JT47 UV', 'MT1083 UV', ...
     'HBR232 UV', ...
     'IB16 UV'}, ...
    {'N2', ...
     'strd-1', 'strd-1', ...
     'cla-1', 'cla-1', 'cla-1', 'cla-1', 'cla-1', ...
     'egl-8', 'egl-8', 'egl-8', 'egl-8', 'egl-8', ...
     'aptf-1', ...
     'ceh-17'} ...
);

% === Appearance ===
defaultColor   = [0.7 0.7 0.7];
baseMarkerSize = 35;
geneMarkerSize = 70;

% === Legend containers ===
legendEntries = [];
legendLabels  = {};

% Painters keeps every element (errorbars, dots) as true vector paths on
% export; without this MATLAB can silently fall back to a rasterized
% OpenGL snapshot for figures with many plotted objects, like this one.
figure('Color','w','Renderer','painters'); hold on; box on;

% ============================================================
%   STRAIN ORDER: plot non-N2 first, N2 last
% ============================================================
strainList = categories(Strain_names_all);

isN2 = false(size(strainList));

for i = 1:numel(strainList)
    sName = strainList{i};

    if isKey(strainToGene, sName) && strcmp(strainToGene(sName), 'N2')
        isN2(i) = true;
    end
end

% N2 first → plotted first → appears underneath
strainList = [strainList(isN2); strainList(~isN2)];

% === legend containers ===
legendEntries = gobjects(0);
legendLabels  = {};

% ellipse smoothness
theta = linspace(0, 2*pi, 120);

% ============================================================
%   LOOP — one SEM ellipse per wormmotel trial directory
% ============================================================

N_each_trial = [];
for s = 1:numel(strainList)

    strainName = strainList{s};

    % all worms for this strain
    idxStrain = Strain_names_all == strainName;

    x_all   = preUV_Qf_all(idxStrain);
    y_all   = postUV_Qf_all(idxStrain);
    dir_all = trialDir_all(idxStrain);

    if isempty(x_all)
        continue;
    end

    % gene mapping
    if isKey(strainToGene, strainName)
        geneName  = strainToGene(strainName);
        geneColor = geneColors(geneName);
    else
        continue;
    end

    % unique wormmotel trial folders for this strain
    uniqueDirs = categories(removecats(dir_all));

    for k = 1:numel(uniqueDirs)

        thisDir = categorical(uniqueDirs(k));
        idxDir  = dir_all == thisDir;

        xDir = x_all(idxDir);
        yDir = y_all(idxDir);

        valid = ~isnan(xDir) & ~isnan(yDir);
        xDir = xDir(valid);
        yDir = yDir(valid);

        if isempty(xDir)
            continue;
        end

        % mean per trial
        xMean = mean(xDir);
        yMean = mean(yDir);

        % SEM per trial
        if numel(xDir) > 1
            N_each_trial(end+1) = numel(xDir);
            xSEM = std(xDir) / sqrt(numel(xDir));
        else
            xSEM = 0;
        end

        if numel(yDir) > 1
            ySEM = std(yDir) / sqrt(numel(yDir));
        else
            ySEM = 0;
        end

        % --- vertical errorbar ---
        h = errorbar(xMean, yMean, ySEM, ...
            'Color', geneColor, ...
            'LineWidth', 1.5, ...
            'CapSize', 0);

        % --- horizontal errorbar ---
        errorbar(xMean, yMean, xSEM, ...
            'horizontal', ...
            'Color', geneColor, ...
            'LineWidth', 1.5, ...
            'CapSize', 0);

        % small center point
        plot(xMean, yMean, '.', ...
            'Color', geneColor, ...
            'MarkerSize', 14);

        % add legend once per gene
        if ~any(strcmp(legendLabels, geneName))
            legendEntries(end+1) = h;
            legendLabels{end+1}  = geneName;
        end
    end
end

fprintf('N max = %d\n', max(N_each_trial));
fprintf('N min = %d\n', min(N_each_trial));
fprintf('N trial = %d\n', size(N_each_trial, 2));

% ============================================================
%   AXES / DIAGONAL / STYLE
% ============================================================
allVals = [preUV_Qf_all(:); postUV_Qf_all(:)];
allVals = allVals(~isnan(allVals));

lims = [min(allVals), max(allVals)];

plot(lims, lims, 'k--', 'LineWidth', 2);

xlim([0 0.4]);
ylim(lims);

xlabel('Pre-UV quiescence fraction');
ylabel('Post-UV quiescence fraction');

set(gca, ...
    'FontSize', 11, ...
    'LineWidth', 1, ...
    'TickDir', 'in', ...
    'Layer', 'top');

grid off;

f = gcf;
f.Units = 'pixels';
f.Position = [100 100 800 800];
pbaspect([1 1 1]);


ax = gca; ax.FontSize = 25;
ax.LineWidth = 2;

hold off;

box off;
% ContentType 'vector' prevents exportgraphics from rasterizing this axes.
exportgraphics(gca, "Fig3B.svg", 'ContentType', 'vector');

%% === Fig3 per-gene panels: trace heatmap + scatter ===
% Outputs:  Fig3C.svg, Fig3E.svg, Fig3G.svg (heatmaps)
%           Fig3D.svg, Fig3F.svg, Fig3H.svg (scatterplots)
clear; clc; close all;

addpath(fullfile('..', 'helpers'));

%% ============================================================
%   STRAIN → GENE MAP
%% ============================================================

strainToGene = containers.Map( ...
    {'N2 UV', ...
     'RB1771 UV', 'RB1775 UV', ...
     'VC387 UV','RB778 UV','RB1777 UV','VC596 UV','VC631 UV', ...
     'CB6614 UV','RM2221 UV','IK777 UV','JT47 UV','MT1083 UV'}, ...
    {'N2', ...
     'strd-1', 'strd-1', ...
     'cla-1','cla-1','cla-1','cla-1','cla-1', ...
     'egl-8','egl-8','egl-8','egl-8','egl-8'} ...
     );

geneList = {'strd-1', 'cla-1', 'egl-8'};
genePanelHeatmap = {'Fig3C', 'Fig3E', 'Fig3G'};   % heatmap panel letter for each gene
genePanelScatter = {'Fig3D', 'Fig3F', 'Fig3H'};   % scatterplot panel letter for each gene

alpha_pre  = 0.05; %#ok<NASGU>
alpha_post = 0.05; %#ok<NASGU>

%% =========================
% CGC-based display labels
%% =========================
strainDisplay = containers.Map( ...
    {'N2 UV', ...
     'RB1771 UV', 'RB1775 UV', ...
     'VC387 UV', 'RB778 UV', 'RB1777 UV', 'VC596 UV', 'VC631 UV', ...
     'CB6614 UV', 'RM2221 UV', 'IK777 UV', 'JT47 UV', 'MT1083 UV'}, ...
    {'N2', ...
     'strd-1(ok2275)', 'strd-1(ok2283)', ...
     'cla-1(ok618)', 'cla-1(ok560)', 'cla-1(ok2285)', 'cla-1(gk352)', 'cla-1(ok937)', ...
     'egl-8(e2917)', 'egl-8(md1971)', 'egl-8(nj77)', 'egl-8(sa47)', 'egl-8(n488)'} ...
    );

%% ============================================================
%   LOAD PRE-FILTERED DATA
%% ============================================================
% data_per_gene.mat is produced by preprocess_per_gene_data.m, which
% scans the raw recording folders once and saves only the rows used by
% this figure (strd-1, cla-1, egl-8 alleles + same-date N2). Re-run
% preprocess_per_gene_data.m if the underlying raw data changes.

load('data_per_gene.mat', 'resultsTable', 'traceTable');

%% ============================================================
%   PER-GENE GROUP ANALYSIS
%% ============================================================

for g = 1:numel(geneList)

    gene = geneList{g};

    %% Find strains for this gene
    strainCats = categories(resultsTable.Strain);
    geneStrains = strings(0,1);

    for i = 1:numel(strainCats)
        s = string(strainCats{i});
        if isKey(strainToGene,s) && strcmp(strainToGene(s),gene)
            geneStrains(end+1,1) = s; %#ok<AGROW>
        end
    end

    geneStrains = unique(geneStrains,'stable');

    strainsToKeep = [geneStrains; "N2 UV"];
    strains_cat = categorical(strainsToKeep, categories(resultsTable.Strain));
    subT = resultsTable(ismember(resultsTable.Strain,strains_cat), :);

    strains_cat_trace = categorical(strainsToKeep, categories(traceTable.Strain));
    subTrace = traceTable(ismember(traceTable.Strain, strains_cat_trace), :);

    %% Same-date control
    mutDates = unique(subT.Date(subT.Strain ~= "N2 UV"));
    subT = subT(subT.Strain ~= "N2 UV" | ismember(subT.Date, mutDates), :);

    mutDates_trace = unique(subTrace.Date(subTrace.Strain ~= "N2 UV"));
    subTrace = subTrace(subTrace.Strain ~= "N2 UV" | ismember(subTrace.Date, mutDates_trace), :);

    %% =========================================================
    %   TRACE HEATMAP FOR THIS GENE GROUP
    %   N2 on top, remaining groups by ascending allele name
    %% =========================================================

    if isempty(subTrace)
        warning('subTrace is empty for gene %s', gene);
        continue;
    end

    % Build display labels for each row
    rowStrains  = string(subTrace.Strain);
    rowAlleles  = strings(size(rowStrains));
    rowIsN2     = false(size(rowStrains));

    for ii = 1:numel(rowStrains)
        s = rowStrains(ii);
        rowIsN2(ii) = (s == "N2 UV");

        if isKey(strainDisplay, s)
            rowAlleles(ii) = strainDisplay(s);
        else
            rowAlleles(ii) = s;
        end
    end

    subTrace.isN2Top = ~rowIsN2;   % N2 -> 0, others -> 1
    subTrace.Allele  = rowAlleles;

    % N2 first, then ascending alphabetical allele, then date
    subTrace = sortrows(subTrace, {'isN2Top','Allele','Date'});

    preCells  = subTrace.datPRE;
    postCells = subTrace.datPOST;

    % Force all traces to row vectors
    preCells  = cellfun(@(x) x(:)', preCells,  'UniformOutput', false);
    postCells = cellfun(@(x) x(:)', postCells, 'UniformOutput', false);

    lenPre  = cellfun(@numel, preCells);
    lenPost = cellfun(@numel, postCells);

    % Keep all nonempty worms only
    keepIdx = (lenPre > 0) & (lenPost > 0);
    traceKeep = subTrace(keepIdx,:);

    preCells  = preCells(keepIdx);
    postCells = postCells(keepIdx);

    if isempty(traceKeep)
        warning('No nonempty traces for gene %s', gene);
        continue;
    end

    % Preserve sorted group order
    alleleOrder = unique(string(traceKeep.Allele), 'stable');
    traceKeep.Allele = categorical(string(traceKeep.Allele), alleleOrder);

    [GtraceKeep, sTraceKeep] = findgroups(traceKeep.Allele);
    nPerStrain = splitapply(@numel, traceKeep.Allele, GtraceKeep);

    edges = cumsum(nPerStrain);
    if ~isempty(edges)
        edges(end) = [];
    end

    % Pad PRE to max PRE length
    maxPreLen = max(cellfun(@numel, preCells));
    alldatPRE_gene = nan(numel(preCells), maxPreLen);

    for ii = 1:numel(preCells)
        x = preCells{ii};
        alldatPRE_gene(ii,1:numel(x)) = x;
    end

    % Pad POST to max POST length
    maxPostLen = max(cellfun(@numel, postCells));
    alldatPOST_gene = nan(numel(postCells), maxPostLen);

    for ii = 1:numel(postCells)
        x = postCells{ii};
        alldatPOST_gene(ii,1:numel(x)) = x;
    end

    [qPRE,~,~]  = fracQ(alldatPRE_gene',1,10,60);
    [qPOST,~,~] = fracQ(alldatPOST_gene',1,10,60);

    % Safe indexing
    startPre  = min(1, size(qPRE,1));
    startPost = min(1, size(qPOST,1));

    qALL = [qPRE(startPre:10:end,:); qPOST(startPost:10:end,:)];

    if isempty(qALL)
        warning('qALL is empty for gene %s', gene);
        continue;
    end

    figure('Color','w','WindowState','maximized');
    imagesc(qALL');

    ax = gca;
    ax.FontSize = 12;
    colorbar
    colormap jet

    starts  = [1; edges(:) + 1];
    stops   = [edges(:); sum(nPerStrain)];
    centers = (starts + stops) / 2;

    y_tick_labels = cellstr(string(sTraceKeep));

    % Italicize mutant allele labels but leave N2 plain
    for k = 1:numel(y_tick_labels)
        if ~strcmp(y_tick_labels{k}, 'N2')
            y_tick_labels{k} = ['\it{' y_tick_labels{k} '}'];
        end
    end

    ax.TickLabelInterpreter = 'tex';
    set(ax, 'YTick', centers, ...
            'YTickLabel', y_tick_labels);

    % X axis in hours
    xticks_steps = 0:360:size(qALL, 1);
    set(gca, 'XTick', xticks_steps + 1);
    set(gca, 'XTickLabel', string(xticks_steps/360));

    hold on
    for k2 = edges(:)'
        plot(xlim, (k2 + 0.5)*[1 1], 'Color', [1 1 1], 'LineWidth', 5);
    end
    hold off

    xlabel('Time (hour)');

    f = gcf;
    f.Units = 'pixels';
    ratio = 0.7;
    f_width = 800/ratio;
    f.Position = [100 100 f_width 800];

    ax = gca;
    ax.FontSize = 29;
    pbaspect([1 ratio 1]);

    exportgraphics(gca, [genePanelHeatmap{g}, '.svg']);

    %% =========================================================
    %   Per-strain stats for scatter plot
    %% =========================================================

    % Group statistics by raw strain first
    [G, sNames] = findgroups(subT.Strain);

    meanPost = splitapply(@mean, subT.Qf_post, G);
    semPost  = splitapply(@(x) std(x)/sqrt(numel(x)), subT.Qf_post, G);
    meanPre  = splitapply(@mean, subT.Qf_pre, G);
    semPre   = splitapply(@(x) std(x)/sqrt(numel(x)), subT.Qf_pre, G);
    nWorms   = splitapply(@numel, subT.Qf_post, G);

    % Pre + Post P-values
    pPost = nan(size(sNames));
    pPre  = nan(size(sNames));

    hPost = nan(size(sNames));
    hPre  = nan(size(sNames));


    disp (gene);
    for i = 1:numel(sNames)

        if sNames(i) == "N2 UV", continue; end

        idxS = subT.Strain == sNames(i);
        dates_i = unique(subT.Date(idxS));

        idxN2 = subT.Strain=="N2 UV";

        wt_post = subT.Qf_post(idxN2);
        wt_pre  = subT.Qf_pre(idxN2);
        dat_post = subT.Qf_post(idxS);
        dat_pre  = subT.Qf_pre(idxS);

        % Rank sum
        alpha_thresh = 0.01;
        tail_type = '';

        if strcmpi(gene,'strd-1')
            tail_type = 'right';
        else
            tail_type = 'left';
        end

        disp(sNames(i))

        [pPre(i), hPre(i)]  = ranksum(wt_pre, dat_pre, 'tail', tail_type, 'alpha', alpha_thresh);
        [pPost(i), hPost(i)] = ranksum(wt_post, dat_post, 'tail', tail_type, 'alpha', alpha_thresh);


        fprintf('median wt_pre = %.2f\n', median(wt_pre));
        fprintf('median dat_pre = %.2f\n', median(dat_pre));
        fprintf('median wt_post = %.2f\n', median(wt_post));
        fprintf('median dat_post = %.2f\n', median(dat_post));

    end

    disp('pPre = ');
    disp(pPre);
    disp('\n\n');

    disp('pPost = ');
    disp(pPost);
    disp('\n\n');

    disp('hPre = ');
    disp(hPre);
    disp('\n\n');

    disp('hPost = ');
    disp(hPost);
    disp('\n\n');

    qPost = pPost;
    qPre = pPre;

    % Build display labels and sort:
    % N2 first, remaining groups by ascending allele name
    displaySortNames = strings(numel(sNames),1);
    isN2_group = false(numel(sNames),1);

    for ii = 1:numel(sNames)
        s = string(sNames(ii));
        isN2_group(ii) = (s == "N2 UV");

        if isKey(strainDisplay, s)
            displaySortNames(ii) = strainDisplay(s);
        else
            displaySortNames(ii) = s;
        end
    end

    mutantIdx = find(~isN2_group);
    [~, mutantLocalOrder] = sort(lower(displaySortNames(mutantIdx)));
    order = [find(isN2_group); mutantIdx(mutantLocalOrder)];

    sNames            = sNames(order);
    meanPost          = meanPost(order);
    semPost           = semPost(order);
    meanPre           = meanPre(order);
    semPre            = semPre(order);
    nWorms            = nWorms(order);
    qPost             = qPost(order);
    qPre              = qPre(order);
    displaySortNames  = displaySortNames(order);

    fprintf('\nGene group: %s\n', gene);
    fprintf('Sample size by allele:\n');
    for ii = 1:numel(displaySortNames)
        fprintf('  %s: n = %d\n', displaySortNames(ii), nWorms(ii));
    end

    %% =========================================================
    %   Scatterplot: N2 first, mutants alphabetical by allele
    %% =========================================================

    close all
    figure('Color','w','WindowState','maximized'); hold on;

    cOrder = colororder;
    Color_preUV  = [0 0 0];
    Color_postUV = cOrder(4,:);

    yl = [0 1];

    % Make sure subT.Strain uses the same order as sNames
    strainCat = categorical(string(subT.Strain), string(sNames));
    groupIdx = grp2idx(strainCat);   % 1..numel(sNames)

    % Build one pooled x/y input for a single beeswarm call
    x_pre  = 2*groupIdx - 1;
    x_post = 2*groupIdx;
    x_all  = [x_pre; x_post];

    y_all  = [subT.Qf_pre; subT.Qf_post];

    % Tick positions
    x_mid = 2*(1:numel(sNames)) - 0.5;

    % One color per unique x position
    % x=1,3,5,... -> pre color
    % x=2,4,6,... -> post color
    groupColors = zeros(2*numel(sNames), 3);
    groupColors(1:2:end, :) = repmat(Color_preUV,  numel(sNames), 1);
    groupColors(2:2:end, :) = repmat(Color_postUV, numel(sNames), 1);

    % Single beeswarm call
    % odd categories = pre-UV (hollow triangle), even categories = post-UV (filled circle)
    mkrs = repmat({'^','o'}, 1, numel(sNames));
    mkrFill = repmat({false, true}, 1, numel(sNames));

    beeswarm(x_all, y_all, ...
        'corral_style', 'random', ...
        'overlay_style', 'sd', ...
        'use_current_axes', true, ...
        'colormap', groupColors, ...
        'dot_size', 3, ...
        'Marker', mkrs, ...
        'MarkerFill', mkrFill, ...
        'MarkerLineWidth', 1.5, ...
        'ChannelWidth', 0.7, ...
        'MarkerFaceColor', '', ...      % leave empty so colormap is used
        'MarkerFaceAlpha', 0.5, ...
        'MarkerEdgeColor', 'w');

    ylabel('Quiescence fraction');


    y_tick_labels = cellstr(displaySortNames);
    % Italicize mutant allele labels but leave N2 plain
    for k = 1:numel(y_tick_labels)
        if ~strcmp(y_tick_labels{k}, 'N2')
            y_tick_labels{k} = ['\it{' y_tick_labels{k} '}'];
        end
    end


    ax = gca;
    ax.TickLabelInterpreter = 'tex';
    set(ax, 'XTick', x_mid, 'XTickLabel', y_tick_labels);
    xtickangle(20)

    box off;

    ylim([yl(1) yl(2)]);
    xlim([0 max(x_post)+1]);

    ylabel('Quiescence fraction');

    x_mid = (x_pre + x_post)/2;

    for i = 1:numel(sNames)
        % text(x_mid(i), yl(1)-0.08, sprintf('n=%d', nWorms(i)), ...
        %     'HorizontalAlignment','center', 'FontSize',12);
    end

    for i = 1:numel(sNames)

        if ~isnan(qPre(i))
            labelPre = sprintf('qpre=%.3f', qPre(i));
            if qPre(i) < 0.05 && sNames(i) ~= "N2 UV"
                labelPre = [labelPre ' *'];
            end
        end

        if ~isnan(qPost(i))
            labelPost = sprintf('qpost=%.3f', qPost(i));
            if qPost(i) < 0.05 && sNames(i) ~= "N2 UV"
                labelPost = [labelPost ' *'];
            end
        end
    end

    box off;

    f = gcf;
    f.Units = 'pixels';

    ratio = 0.55;
    f_width = 800/ratio;
    f.Position = [100 100 f_width 800];

    ax = gca;
    ax.FontSize = 34;
    ax.LineWidth = 2;
    pbaspect([1 ratio 1]);

    exportgraphics(gca, [genePanelScatter{g}, '.svg']);

end

%% === Local functions (SKAT percentile lookup used in the Fig3A block) ===

function pct = lookupSkatPercentile(genotypeStr, byCommon, bySetid, aliasMap)
%LOOKUPSKATPERCENTILE Match gene name(s) parsed from a Genotype string to
%the SKAT percentile rank, taking the lowest (most significant) value
%across loci for multi-gene genotypes. Returns NaN if no gene matches.
    genes = extractGenesFromGenotype(genotypeStr);
    pcts = [];
    for i = 1:numel(genes)
        candidates = {genes{i}};
        if isKey(aliasMap, genes{i})
            candidates{end+1} = aliasMap(genes{i}); %#ok<AGROW>
        end
        for c = 1:numel(candidates)
            g = candidates{c};
            if isKey(byCommon, g)
                pcts(end+1) = byCommon(g); %#ok<AGROW>
                break;
            elseif isKey(bySetid, g)
                pcts(end+1) = bySetid(g); %#ok<AGROW>
                break;
            end
        end
    end
    if isempty(pcts)
        pct = NaN;
    else
        pct = min(pcts);
    end
end

function ids = resolveGeneID(genotypeStr, commonToSetid, setidToSetid, aliasMap)
%RESOLVEGENEID Canonicalize each gene parsed from a Genotype string to its
%SKAT SetID (a gene can appear in the tsv under Common_name or SetID), so
%different strains/alleles of the same gene collapse to one ID. Returns a
%string array (possibly >1 entry for multi-locus genotypes; empty if no
%gene in the string has a SKAT match).
    genes = extractGenesFromGenotype(genotypeStr);
    ids = strings(0,1);
    for i = 1:numel(genes)
        candidates = {genes{i}};
        if isKey(aliasMap, genes{i})
            candidates{end+1} = aliasMap(genes{i}); %#ok<AGROW>
        end
        for c = 1:numel(candidates)
            g = candidates{c};
            if isKey(commonToSetid, g)
                ids(end+1,1) = string(commonToSetid(g)); %#ok<AGROW>
                break;
            elseif isKey(setidToSetid, g)
                ids(end+1,1) = string(setidToSetid(g)); %#ok<AGROW>
                break;
            end
        end
    end
end

function pctList = lookupSkatPercentiles(genotypeStr, byCommon, bySetid, aliasMap)
%LOOKUPSKATPERCENTILES Like lookupSkatPercentile, but returns one
%percentile per locus in genotype order (NaN for a locus with no SKAT
%match) instead of collapsing multi-gene genotypes to the minimum.
    genes = extractGenesFromGenotype(genotypeStr);
    pctList = nan(1, numel(genes));
    for i = 1:numel(genes)
        candidates = {genes{i}};
        if isKey(aliasMap, genes{i})
            candidates{end+1} = aliasMap(genes{i}); %#ok<AGROW>
        end
        for c = 1:numel(candidates)
            g = candidates{c};
            if isKey(byCommon, g)
                pctList(i) = byCommon(g);
                break;
            elseif isKey(bySetid, g)
                pctList(i) = bySetid(g);
                break;
            end
        end
    end
end

function genes = extractGenesFromGenotype(genotypeStr)
%EXTRACTGENESFROMGENOTYPE Pull the gene identifier(s) preceding "(allele)"
%for each ';'-separated locus in a genotype string, e.g.
%"gei-1(gk3062) III; C25A8.5(gk1224) IV." -> {'gei-1','C25A8.5'}.
    genes = {};
    loci = strsplit(genotypeStr, ';');
    for i = 1:numel(loci)
        tok = regexp(strtrim(loci{i}), '^([^()]+)\(', 'tokens', 'once');
        if isempty(tok)
            continue;
        end
        parts = strsplit(strtrim(tok{1}), {'/', ','});
        for p = 1:numel(parts)
            g = strtrim(parts{p});
            if ~isempty(g)
                genes{end+1} = g; %#ok<AGROW>
            end
        end
    end
end
```
