## Supplementary material for "A semi-automated genetic screen using genome-wide association analysis identifies modulators of stress-induced sleep in *Caenorhabditis elegans*": Software S1: preprocess_per_gene_data.m

```
%% One-time preprocessing for Figure 3 per-gene panels.
% Scans the raw recording folders on disk, builds the per-worm tables used
% by the heatmap + scatter panels in Main_figure3.m, filters them down to
% the strains of interest (strd-1, cla-1, egl-8 alleles + same-date N2),
% and saves a lightweight data_per_gene.mat next to this script.
%
% You only need to run this once, on a machine that has the raw recordings.
% The resulting data_per_gene.mat is what gets committed to the repo and
% loaded by Main_figure3.m at publication time.

clear; clc;

%% ============================================================
%   PATHS  --  set these to your local raw-recording folders
%% ============================================================

baseDirs = { ...
    'C:\Users\jl200\Dropbox\Shared_folders\Helya_John_Shared_folder\Writing\Raw_data\SleepScreenSelectedMOMO', ...
    'C:\Users\jl200\Dropbox\Shared_folders\Helya_John_Shared_folder\Writing\Raw_data\SleepScreenSelectedKAWKAB'};

outFile = fullfile(fileparts(mfilename('fullpath')), 'data_per_gene.mat');

%% ============================================================
%   STRAIN -> GENE MAP (strains we keep; everything else is dropped)
%% ============================================================

strainToGene = containers.Map( ...
    {'N2 UV', ...
     'RB1771 UV', 'RB1775 UV', ...
     'VC387 UV','RB778 UV','RB1777 UV','VC596 UV','VC631 UV', ...
     'CB6614 UV','RM2221 UV','IK777 UV','JT47 UV','MT1083 UV'}, ...
    {'N2', ...
     'strd-1', 'strd-1', ...
     'cla-1','cla-1','cla-1','cla-1','cla-1', ...
     'egl-8','egl-8','egl-8','egl-8','egl-8'} ...
     );

%% ============================================================
%   LOAD DATA
%% ============================================================

allStrains = strings(0,1);
allPost    = [];
allPre     = [];
allDates   = strings(0,1);
alldatPRE  = {};
alldatPOST = {};

for b = 1:numel(baseDirs)

    root = baseDirs{b};
    folders = dir(root);
    folders = folders([folders.isdir] & ~ismember({folders.name},{'.','..'}));

    for i = 1:numel(folders)

        sub = folders(i).name;
        matFile = fullfile(root,sub,'Analysis',[sub '_ForPlot.mat']);
        if ~exist(matFile,'file'), continue; end

        S = load(matFile,'Qf_output_struct');

        analysisDir = fullfile(root, sub, 'Analysis');

        FilePRE  = dir(fullfile(analysisDir, '*_NS.mat'));
        FilePOST = dir(fullfile(analysisDir, '*_UVC.mat'));

        if isempty(FilePRE) || isempty(FilePOST)
            continue;
        end

        Spre   = load(fullfile(analysisDir, FilePRE(1).name));
        datPRE = Spre.ActVal;

        Spost   = load(fullfile(analysisDir, FilePOST(1).name));
        datPOST = Spost.ActVal;

        strains = string(S.Qf_output_struct.strain_category);
        strains = regexprep(strtrim(strains),'^D\d+-','');

        post = S.Qf_output_struct.Qf_4hr_postUV(:);
        pre  = S.Qf_output_struct.Qf_30min_preUV(:);

        n = min([numel(post), numel(pre), size(datPRE,1), size(datPOST,1), numel(strains)]);

        strains = strains(1:n);
        post    = post(1:n);
        pre     = pre(1:n);
        datPRE  = datPRE(1:n,:);
        datPOST = datPOST(1:n,:);

        dateToken = string(regexp(sub,'^\d{2}-\d{2}-\d+','match','once'));

        allStrains = [allStrains; strains];
        allPost    = [allPost; post];
        allPre     = [allPre; pre];
        allDates   = [allDates; repmat(dateToken,n,1)];

        for k = 1:n
            alldatPRE{end+1,1}  = datPRE(k,:);   %#ok<AGROW>
            alldatPOST{end+1,1} = datPOST(k,:);  %#ok<AGROW>
        end
    end
end

resultsTable = table( ...
    categorical(allStrains), ...
    allPost, allPre, categorical(allDates), ...
    'VariableNames',{'Strain','Qf_post','Qf_pre','Date'});

traceTable = table( ...
    categorical(allStrains), ...
    categorical(allDates), ...
    alldatPRE(:), ...
    alldatPOST(:), ...
    'VariableNames',{'Strain','Date','datPRE','datPOST'});

%% ============================================================
%   FILTER TO GENE STRAINS + SAME-DATE N2
%% ============================================================

allowedStrains = string(keys(strainToGene));
allowedCat = categorical(allowedStrains, categories(resultsTable.Strain));

resultsTable = resultsTable(ismember(resultsTable.Strain, allowedCat), :);

mutDates = unique(resultsTable.Date(resultsTable.Strain ~= "N2 UV"));
resultsTable = resultsTable( ...
    resultsTable.Strain ~= "N2 UV" | ...
    (resultsTable.Strain=="N2 UV" & ismember(resultsTable.Date,mutDates)), :);

allowedCat_trace = categorical(allowedStrains, categories(traceTable.Strain));
traceTable = traceTable(ismember(traceTable.Strain, allowedCat_trace), :);

mutDates_trace = unique(traceTable.Date(traceTable.Strain ~= "N2 UV"));
traceTable = traceTable( ...
    traceTable.Strain ~= "N2 UV" | ...
    (traceTable.Strain=="N2 UV" & ismember(traceTable.Date,mutDates_trace)), :);

%% ============================================================
%   SAVE
%% ============================================================

save(outFile, 'resultsTable', 'traceTable', '-v7.3');
fprintf('Saved %s\n', outFile);
fprintf('  resultsTable: %d rows\n', height(resultsTable));
fprintf('  traceTable:   %d rows\n', height(traceTable));
```
