## Supplementary material for "A semi-automated genetic screen using genome-wide association analysis identifies modulators of stress-induced sleep in *Caenorhabditis elegans*": Software S1: beeswarm.m

```
function x = beeswarm(x,y,varargin)
%function xbee = beeswarm(x,y)
%
% Input arguments:
%   x               column vector of groups (only tested for integer)
%   y               column vector of data
%
% Optional input arguments:
%   sort_style      ('nosort' - default | 'up' | 'down' | 'fan' | 'rand' | 'square' | 'hex')
%   corral_style    ('none' default | 'gutter' | 'omit' | 'rand')
%   dot_size        relative. default=1
%   overlay_style   (false default | 'box' | 'sd' | 'ci')
%   use_current_axes (false default | true)
%   colormap        (lines default | 'jet' | 'parula' | 'r' | Nx3 matrix of RGB values]
%
% Output arguments:
%   xbee            optimized layout positions
%
% Known Issues:
%       x locations depend on figure aspect ratio. resizing the figure window and rerunning may give different results
%       setting corral to 'none' still has a gutter when the width is large
%
% Usage example:
% 	x = round(rand(150,1)*5);
%   y = randn(150,1);
%   beeswarm(x,y,3,'sort_style','up','overlay_style','ci')
%
% % Ian Stevenson, CC-BY 2019

p = inputParser;
addRequired(p,'x')
addRequired(p,'y')
validScalarPosNum = @(x) isnumeric(x) && isscalar(x) && (x > 0);
addOptional(p,'sort_style','nosort')
addOptional(p,'corral_style','none')
addOptional(p,'dot_size',11/sqrt(length(x)),validScalarPosNum)
addOptional(p,'overlay_style',false)
addOptional(p,'use_current_axes',false)
addOptional(p,'colormap','lines')
addOptional(p,'MarkerFaceColor','')
addOptional(p,'MarkerFaceAlpha',.5)
addOptional(p,'MarkerEdgeColor','none')
addOptional(p,'Marker','o') % single marker, or cell array (one per category)
addOptional(p,'MarkerFill',true) % single logical, or cell array (one per category)
addOptional(p,'MarkerLineWidth',0.5) % outline thickness for hollow markers; single value, or cell array (one per category)
addOptional(p,'ChannelWidth',.9,validScalarPosNum) % fraction of the unit gap between categories that the swarm may use; lower it to leave more empty space between adjacent groups
parse(p,x,y,varargin{:});

% extra parameters
rwid = .05; % width of overlay box/dash

dcut=8; % spacing factor
nxloc=512; % resolution for optimization
chanwid = p.Results.ChannelWidth; % percent width of channel to use
yl = [min(y) max(y)]; % default y-limits
asp_rat = 1;
keep_hold = false;

% get aspect ratio for a figure window
if isfinite(p.Results.dot_size)
    if ~p.Results.use_current_axes
        % make new axes
        s=scatter(x,y);
        xl=[min(x)-.5 max(x)+.5];
    else
        xl=xlim();
    end
    yl=ylim();
    pasp_rat = get(gca,'PlotBoxAspectRatio');
    dasp_rat = get(gca,'DataAspectRatio');
    asp_rat = pasp_rat(1)/pasp_rat(2);

    % pix-scale
    pf = get(gcf,'Position');
    pa = get(gca,'Position');
    as = pf(3:4).*pa(3:4); % width and height of panel in pixels
    dcut = dcut*sqrt(p.Results.dot_size)/as(1)*(range(unique(x))+1);
    if ~ishold
        cla
    else
        keep_hold = true;
    end
end

% sort/round y for different plot styles
yorig=y;
switch lower(p.Results.sort_style)
    case 'up'
        [y,sid]=sort(y);
    case 'fan'
        [~,sid]=sort(abs(y-mean(y)));
        sid=[sid(1:2:end); sid(2:2:end)];
        y=y(sid);
    case 'down'
        [y,sid]=sort(y,'descend');
    case 'rand'
        sid=randperm(length(y));
        y=y(sid);
    case 'square'
        nxloc=.9/dcut;
%         [~,e,b]=histcounts(y,ceil((range(x)+1)*chanwid*nxloc/2/asp_rat));
        edges = linspace(min(yl),max(yl),ceil((range(x)+1)*chanwid*nxloc/asp_rat));
        [~,e,b]=histcounts(y,edges);
        y=e(b)'+mean(diff(e))/2;
        [y,sid]=sort(y);
    case 'hex'
        nxloc=.9/dcut;
%         [~,e,b]=histcounts(y,ceil((range(x)+1)*chanwid*nxloc/2/sqrt(1-.5.^2)/asp_rat));
        edges = linspace(min(yl),max(yl),ceil((range(x)+1)*chanwid*nxloc/sqrt(1-.5.^2)/asp_rat));
        [n,e,b]=histcounts(y,edges);
        oddmaj=0;
        if sum(mod(n(1:2:end),2)==1)>sum(mod(n(2:2:end),2)==1),
            oddmaj=1;
        end
        y=e(b)'+mean(diff(e))/2;
        [y,sid]=sort(y);
        b=b(sid);
    otherwise
        sid=1:length(y);
end
x=x(sid);
yorig=yorig(sid);
[ux,~,ic] = unique(x);
% rmult=(range(ux)+1)*2;
rmult=5;

% for each group...
for i=1:length(ux)
    fid = find(ic==i);

    % set of possible x locations
    xi = linspace(-chanwid/2*rmult,chanwid/2*rmult,nxloc*rmult+(mod(nxloc*rmult,2)==0))'+ux(i);

    % rescale y to that things are square visually
    zy=(y(fid)-min(yl))/(max(yl)-min(yl))/asp_rat*(range(ux)+1)*chanwid;

    % precalculate y distances so that we only worry about nearby points
    D0=squareform(pdist(zy))<dcut*2;

    if length(fid)>1
        % for each data point in the group sequentially...
        for j=1:length(fid)
            if strcmp(lower(p.Results.sort_style),'hex')
                xi = linspace(-chanwid/2*rmult,chanwid/2*rmult,nxloc*rmult+(mod(nxloc*rmult,2)==0))'+ux(i);
                if mod(b(fid(j)),2)==oddmaj
                    xi = linspace(-chanwid/2*rmult,chanwid/2*rmult,nxloc*rmult+(mod(nxloc*rmult,2)==0))'+ux(i)+mean(diff(xi))/2;
                end
            end
            zid = D0(j,1:j-1);
            e = (xi-ux(i)).^2; % cost function
            if ~strcmp(lower(p.Results.sort_style),'hex') && ~strcmp(lower(p.Results.sort_style),'square')
                if sum(zid)>0
                    D = pdist2([xi ones(length(xi),1)*zy(j)], [x(fid(zid)) zy(zid)]);
                    D(D<=dcut)=Inf;
                    D(D>dcut & isfinite(D))=0;
                    e = e + sum(D,2) + randn(1)*10e-6; % noise to tie-break
                end
            else
                if sum(zid)>0
                    D = pdist2([xi ones(length(xi),1)*zy(j)], [x(fid(zid)) zy(zid)]);
                    D(D==0)=Inf;
                    D(D>dcut & isfinite(D))=0;
                    e = e + sum(D,2) + randn(1)*10e-6; % noise to tie-break
                end
            end

            if strcmp(lower(p.Results.sort_style),'one')
                e(xi<ux(i))=Inf;
            end
            [~,mini] = min(e);
            if mini==1 && rand(1)>.5, mini=length(xi); end
            x(fid(j)) = xi(mini);
        end
    end
%     x(fid)=x(fid)-median(x(fid))+ux(i); % center x locations by median
end

if strcmp(lower(p.Results.sort_style),'randn')
    x=ux(ic)+randn(size(ic))/4;
end

% corral any points outside of the channel
out_of_range = abs(x-ux(ic))>chanwid/2;
switch lower(p.Results.corral_style)
    case 'gutter'
        id = (x-ux(ic))>chanwid/2;
        x(id)=chanwid/2+ux(ic(id));
        id = (x-ux(ic))<-chanwid/2;
        x(id)=-chanwid/2+ux(ic(id));
    case 'omit'
        x(out_of_range)=NaN;
    case 'random'
        x(out_of_range)=ux(ic(out_of_range))+rand(sum(out_of_range),1)*chanwid-chanwid/2;
end

% plot groups and add overlay
if isfinite(p.Results.dot_size)
    if isnumeric(p.Results.colormap)
        cmap=p.Results.colormap;
    else
        cmap = feval(p.Results.colormap,length(ux));
    end
    for i=1:length(ux)
        if iscell(p.Results.Marker)
            mkr = p.Results.Marker{i};
        else
            mkr = p.Results.Marker;
        end
        if iscell(p.Results.MarkerFill)
            doFill = p.Results.MarkerFill{i};
        else
            doFill = p.Results.MarkerFill;
        end
        if isempty(p.Results.MarkerFaceColor')
            faceColor = cmap(i,:);
        else
            faceColor = p.Results.MarkerFaceColor;
        end
        if iscell(p.Results.MarkerLineWidth)
            lw = p.Results.MarkerLineWidth{i};
        else
            lw = p.Results.MarkerLineWidth;
        end
        if doFill
            scatter(x(ic==i),y(ic==i),p.Results.dot_size*36,'filled','Marker',mkr,'MarkerFaceAlpha', ...
                p.Results.MarkerFaceAlpha,'MarkerEdgeColor',p.Results.MarkerEdgeColor,'MarkerFaceColor',faceColor)
        else
            % hollow marker: color the outline instead of the (nonexistent) fill
            scatter(x(ic==i),y(ic==i),p.Results.dot_size*36,'Marker',mkr,'MarkerEdgeColor',faceColor,'LineWidth',lw)
        end
        hold on
        iqr = prctile(yorig(ic==i),[25 75]);
        switch lower(p.Results.overlay_style)
            case 'box'
                rectangle('Position',[ux(i)-rwid iqr(1) 2*rwid iqr(2)-iqr(1)],'EdgeColor','k','LineWidth',2)
                line([ux(i)-rwid ux(i)+rwid],[1 1]*median(yorig(ic==i)),'LineWidth',3,'Color',cmap(i,:))
            case 'sd'
                % erbar_color = cmap(i,:);
                erbar_color = 'b';
                line([1 1]*ux(i),mean(yorig(ic==i))+[-1 1]*std(yorig(ic==i)),'Color',erbar_color,'LineWidth',4)
                line([ux(i)-4*rwid ux(i)+4*rwid],[1 1]*mean(yorig(ic==i)),'LineWidth',4,'Color',erbar_color)
            case 'se'
                line([1 1]*ux(i),mean(yorig(ic==i))+[-1 1]*std(yorig(ic==i))/sqrt(sum(ic==i)),'Color',cmap(i,:),'LineWidth',4)
                line([ux(i)-2*rwid ux(i)+2*rwid],[1 1]*mean(yorig(ic==i)),'LineWidth',4,'Color',cmap(i,:))
            case 'ci'
                line([1 1]*ux(i),mean(yorig(ic==i))+[-1 1]*std(yorig(ic==i))/sqrt(sum(ic==i))*tinv(0.975,sum(ic==i)-1),'Color',cmap(i,:),'LineWidth',2)
                line([ux(i)-2*rwid ux(i)+2*rwid],[1 1]*mean(yorig(ic==i)),'LineWidth',3,'Color',cmap(i,:))
        end

    end
    hold off
    if keep_hold
        hold on
    end
    xlim(xl)
    ylim(yl)
end

% unsort so that output matches the original y data
x(sid)=x;

end
```
