## Supplementary material for "A semi-automated genetic screen using genome-wide association analysis identifies modulators of stress-induced sleep in *Caenorhabditis elegans*": Software S1: fracQ.m

```
%fracQ.m
% Calculates the fraction of quiescence from activity data in p.
% p is the transpose of the activity trace.
% qt is the number of pixels required to move to call the frame not-quiescent.
%   1 is a good value.
% spi is the number of seconds each image represents.
% ps is the moving average size. ps must be odd. If it is not, the program
%   automatically adds 1 to ps.
%
% fracQ returns PAV, a fraction-of-quiescence vector equal in size to p,
% and t1 which is the time vector associated with PAV.

function [PAV t1 pnew] = fracQ(p,qt,spi,ps)

if mod(ps,2)==0
    ps=ps+1;
end

pq=p<qt;
[sizex sizey]=size(p);
t1=[1:1/spi:sizex]/3600*spi;
rois=1:sizey;
pstart=ceil(ps/2);
pf=floor(ps/2);

pav=zeros(ceil(sizex/ps),sizey);
for i=pstart:ps:sizex-ps
    for j=1:sizey
        pav((i+pf)/ps,j)=mean(pq((i-pf):(i+pf),j), "omitnan");
    end
end
pax=[1:ceil(sizex/ps)]*ps/3600*spi;
pax2=[1:sizex]/3600*spi;
PAV=interp2(rois,pax',pav,rois,t1');
pnew=interp2(rois,pax2',p,rois,t1')/spi;

end
```
