## Supplementary material for "A semi-automated genetic screen using genome-wide association analysis identifies modulators of stress-induced sleep in *Caenorhabditis elegans*": Software S1: fracQbin230817.m

```
%fracQbin function (normalizes post-treatment data to pre-treatment data)
%identifies periods of quiescence based on two criteria:
%for a given epoch to be considered quiescent it must meet the following
%criteria:
%(1) ActVal must be below ACTIVITY THRESHOLD
%(2) fracQ must be above QUIESCENCE THRESHOLD
%in this script the activity threshold is established relative to an
%untreated control group. the activity threshold is 1% of the 95th
%percentile of ActVal for control animals (median of 95th percentiles from each animal)
%fracQ is the fraction of epochs in a 10 minute moving window
%around a given epoch (5 mins before/5mins after), where ActVal falls below
%the activity threshold
%after calculating fracQ, this fxn will sum up the time spent quiescent
%into bins, as well as average quiescence (normalized)

%inputs:
%pre and post (MxN matrices): Activity Matrices w/ #pixels changed in
%subtracted frames (from before/after UV respectively)
%%M (rows) are successive time frames, N (cols) are wells
%%ActVal'

%gp (cell array): each cell contains vector of well numbers pertaining to
%%each group

%ATp (scalar): activity threshold percentage. this is the percentage of the
%%'max' activity in untreated groups that will be used as activity threshold
%%note ATp is a fraction (if you want 1%, then ATp=0.001)

%QT (scalar): quiescence threshold. this is the percentage of time in a
%%moving window where an animal must be inactive to be considered quiescent
%%0.25 is a good staring point

%qWin (scalar): quiescence window. this is the size of the moving average
%%window used to calculate fracQ at each point
%%use number of epochs, eg for 10min window w/ images every 10secs, qWin=60

%binSz (scalar): size of bins (in # of epochs) to add up qui and act into

%EPM (scalar): epochs per minute (eg 1 image per 10sec = 6 epochs per min)

%outputs:
%qTime (cell array): each cell contains matrix of time spent quiescent
%%values are minutes
%%rows are binned time points and cols are wells

%ActNorm (cell array): each cell contains activity values divided by the
%%normalization factor (95th percentile of control activity)
%%rows are time points, cols are wells

%ActAve (cell array): average ActNorm value for each bin

function [qTime,ActNorm,ActAve,qTimePRE,ActAvePRE,durPre] = fracQbin230817(pre,post,gp,ATp,QT,qWin,binSz,EPM)

nGp=length(gp); %get number of groups used for analysis

mRowPRE=size(pre,1); %get size of pre data
mRow=size(post,1); %get size of post data

%if qWin is an even number, add 1
if mod(qWin,2)>0
    qWin=qWin+1;
end

%since we are using a moving window average
%there will be a period of time at the beginning and end of the recording
%that is not covered by a sufficiently large time window to give the data
winLag=floor(qWin/2); %find the lagging window size (number of epochs before or after)

stRanPRE=winLag + 1; %find the first position covered by moving window
enRanPRE=mRowPRE - winLag; %find the last position covered by moving window
RanPRE=[stRanPRE:enRanPRE]; %find all positions covered by moving window
bigRanPRE=[stRanPRE-winLag:enRanPRE+winLag]; %find all positions used for moving mean
durPre=length(RanPRE) * (1/EPM);

stRan=winLag + 1; %find the first position covered by moving window
enRan=mRow - winLag; %find the last position covered by moving window
Ran=[stRan:enRan]; %find all positions covered by moving window
bigRan=[stRan-winLag:enRan+winLag]; %find all positions used for moving mean

%find normalization-factor (NF)
%median of 95th percentile of ActVal in preData for each group
NF=zeros(1,nGp);
for i = 1:nGp
    wellsI=gp{i};
    pct95i=prctile(pre(:,wellsI)',95);
    NF(i)=median(pct95i,'omitnan');
end
clear i gpI wellsI pct95i

%make a matrix for each group that is normalized to its baseline
ActNormPre=cell(1,nGp);
QuiNormPre=cell(1,nGp);
fracQpre=cell(1,nGp);
qOutPre=cell(1,nGp);
ActNorm=cell(1,nGp);
QuiNorm=cell(1,nGp);
fracQ=cell(1,nGp);
qOut=cell(1,nGp);
for j = 1:nGp
    NFj=NF(j); %get the normalization factor for this group
    wellsJ=gp{j}; %get the wells for this group

    %make binary for inactivity based on activity threshold for this group
    ATj = NFj * ATp; %activity threshold specific to this group

    %pre-treatment analysis
    ActJpre=pre(bigRanPRE,wellsJ); %get actval for these wells in the time range
    ActNormPre{j}=ActJpre(RanPRE,:) * (1/NFj); %divide all activity values by norm factor
    QuiNormPre{j}=ActNormPre{j} <= ATp; %find all times where normalized activity is below activity threshold
    QuiJpre=ActJpre;
    QuiJpre=QuiJpre <= ATj; %QuiJ is essentially the same as QuiNorm (all values less than ATp percent of normalization factor)
    fracQjPRE=movmean(QuiJpre,qWin); %fracQ is a moving average of time spent quiescent
    fracQpre{j}=fracQjPRE(RanPRE,:);
    qnJpre=QuiNormPre{j};
    frJpre=fracQpre{j} >= QT;
    qOutPre{j}= qnJpre .* frJpre; %qui = times when: qnJpre=1 (activity below activity threshold); AND frJpre=1 (fracQ above quiescence threshold)

    %post-treatment analysis
    ActJ=post(bigRan,wellsJ); %get actval for these wells in the time range
    ActNorm{j}=ActJ(Ran,:) * (1/NFj); %divide all activity values by norm factor
    QuiNorm{j}=ActNorm{j} <= ATp; %find all times where normalized activity is below activity threshold
    QuiJ=ActJ;
    QuiJ=QuiJ <= ATj; %QuiJ is essentially the same as QuiNorm (all values less than ATp percent of normalization factor)
    fracQj=movmean(QuiJ,qWin); %fracQ is a moving average of time spent quiescent
    fracQ{j}=fracQj(Ran,:); %take only range covered by window size
    qnJ=QuiNorm{j}; %=1 if activity is below ATp
    frJ=fracQ{j} >= QT; %=1 if fracQ is above QT
    qOut{j} = qnJ .* frJ; %multiply (pairwise) so that only QUI if both <ATp and >QT
end
clear j

%sum quiescent time and activity according to binSize
RanLen=length(Ran); %get number of epochs covered by range
Nbins = floor(RanLen/binSz); %find number of bins to cut this range into
binStarts=[1:binSz:Nbins*binSz+1]; %find start times for each bin
binEnds=binStarts + (binSz-1); %find end times for each bin
qTime=cell(1,nGp); %set output variable for total quiescent time
ActAve=cell(1,nGp); %set output variable for activity in each bin
%for pre-treatment, sum up the entire time frame (minus the lag window)
%then transform the time frame so that it equals bin size of post-treatment
RanLenPRE=length(RanPRE);
qTimePRE=cell(1,nGp);
ActAvePRE=cell(1,nGp);
for i = 1:nGp %loop for each group
    quiI=qOut{i}; %get qui data for this group
    actI=ActNorm{i}; %get act data for this group

    %pre
    qTimePRE{i}=sum(qOutPre{i},'omitnan') * (1/EPM); %convert qui time in minutes
    qTimePRE{i}=qTimePRE{i} * (binSz/RanLenPRE); %scale qui-time total to per hour
    ActAvePRE{i}=sum(ActNormPre{i},'omitnan') * (1/RanLenPRE); %average quiescent value over all epochs

    %post
    for j = 1:Nbins
        ranJ=binStarts(j):binEnds(j);
        qTime{i}(j,:)=sum(quiI(ranJ,:),'omitnan') * (1/EPM);
        ActAve{i}(j,:)=sum(actI(ranJ,:),'omitnan') * (1/binSz);
    end

end


end
```
