## Supplementary material for "A semi-automated genetic screen using genome-wide association analysis identifies modulators of stress-induced sleep in *Caenorhabditis elegans*": Software S1: prismXY0915.m

```
%turn matrix from XY data into prism input (eg timecourse from fracQbin)

%cellArray: cell array of MxN matrices, M (subjects), N (time points),
%could be output from fracQbin
%N (scalar): maximum size of group in output
%ord (vector): order to arrange output(correspond to indices of cellArray)
%X (vector): x-values corresponding to rows of matrices in cellArray

%prismOut (matrix): input for prism XY graph, each group is N-members long
%with NaN filling in blank spaces. first column is X-vector

function prismOut=prismXY0915(cellArray,N,ord,X)

nGps=length(ord); %how many groups are in new matrix
nRows=length(X); %how many rows are in new matrix
newMat=NaN(nRows,nGps*N); %make new dummy matrix
for i = 1:nGps
    gpI=ord(i); %get group number for this loop
    matI=cellArray{gpI}; %get matrix for this loop
    [mI,nI]=size(matI);
    colSt=(i-1)*N + 1; %get first column for input
    colEn=colSt + nI - 1; %get last column for input
    newMat(:,colSt:colEn)=matI(1:nRows,:);
end

if nRows==size(X,1) %if X is row vector, transpose it
    prismOut=[X,newMat];
else
    prismOut=[X',newMat];
end

end
```
