## Supplementary material for "A semi-automated genetic screen using genome-wide association analysis identifies modulators of stress-induced sleep in *Caenorhabditis elegans*": Software S1: README.md

# helpers/

Shared MATLAB helper functions used by the figure scripts in this
repository.

## Functions

### `fracQ.m`
Converts a raw activity trace to a **fraction-of-quiescence** trace.
Given a matrix of per-frame pixel-change values, an activity threshold
`qt` (pixels needed to call a frame "not quiescent"), the seconds-per-image
`spi`, and a moving-window size `ps`, it returns the per-window fraction of
quiescent frames and a matched time vector. Used to build the quiescence
heatmaps in Figures 1 and 3.

### `fracQbin230817.m`
Quiescence-detection routine that **normalizes post-treatment data to a
pre-treatment baseline**. A frame is called quiescent when (i) its activity
falls below an activity threshold (defined as a fraction of the
95th-percentile activity of an untreated control group), and (ii) within a
10-minute moving window centered on that frame, the fraction of frames
below the activity threshold exceeds a quiescence threshold. The function
then bins quiescence in time and returns the binned/averaged values used
for the group-level summaries in Figures 1–3.

### `prismXY0915.m`
Data-reshaping utility. Takes a cell array
of per-group MxN matrices (subjects × time points) and concatenates them
into a single matrix with one shared X column followed by `nGroups × N`
data columns, NaN-padded so all groups have the same number of subject
columns. The output layout matches the **XY-table format used by GraphPad
Prism**, which makes it convenient to copy-paste into Prism for external
plotting/statistics, but the function itself is just a MATLAB array
transformation — GraphPad Prism is not a dependency of this repository.

### `beeswarm.m`
Third-party beeswarm/violin-style scatter plotting function (Ian Stevenson,
CC-BY 2019). Computes optimized x-positions for a column of grouped data so
points don't overlap, with options for sorting style, corralling, dot size,
overlay statistics (box / SD / CI), and color maps. Used for the per-strain
scatter plots in Figures 1 and 3.
