## Supplementary material for "A semi-automated genetic screen using genome-wide association analysis identifies modulators of stress-induced sleep in *Caenorhabditis elegans*": Software S1: README.md

# Software S1

Code and data accompanying the manuscript:

"**A semi-automated genetic screen identifies modulators of stress-induced sleep in *Caenorhabditis elegans* through genome-wide association analysis**"
authored by Zihao (John) Li *et al.*, Fang-Yen Laboratory, Department of Biomedical Engineering, The Ohio State University.

---

## Folder layout

```
.
├── figure1/         Figure 1 plotting script + data
├── figure2/         Figure 2 plotting script + data
├── figure3/         Figure 3 plotting script + data
├── helpers/         Shared MATLAB helper functions used by the figure scripts
├── tableS1/         Script to build Supplemental Table S1
├── SKAT/            R/SKAT pipeline used for the association analysis
├── LICENSE          Apache License 2.0
└── README.md
```

## Reproducing figures
Each `figureN/` folder is self-contained. To regenerate paper figures:

1. Open MATLAB, `cd` into the corresponding `figureN/` folder.
2. Run `Main_figureN.m`. Figures display on screen and are saved as SVG files
into the same folder. `Main_figure2.m` and `Main_figure3.m` additionally write out
a supplemental CSV of the per-strain data.

## Reproducing Table S1
[tableS1/](tableS1/) combines the per-strain CSVs written by `Main_figure2.m` and
`Main_figure3.m` (see Reproducing figures above) into Table S1 (`tableS1/Table_S1.xlsx`), with one
sheet for `MMP_screen` and another for `candidate_screen`. To regenerate it:

1. Run `Main_figure2.m` and `Main_figure3.m` (see above) so their CSVs are up to date.
2. Open MATLAB, `cd` into `tableS1/`, and run `Make_Table_S1.m`.
3. `tableS1/Table_S1.xlsx` is written (overwriting any previous copy).

## SKAT analysis
The SKAT analysis pipeline lives in
[SKAT/](SKAT/); see [SKAT/README.md](SKAT/README.md) for how to run it.

## Helper functions
Shared MATLAB helper functions live in [helpers/](helpers/); see
[helpers/README.md](helpers/README.md) for a description of each one.

## Software requirements

**MATLAB** (tested on R2023b or later)
- Statistics and Machine Learning Toolbox (`ranksum`, `vartestn`)

**R** (for the SKAT pipeline only; tested on R 4.3+)
- See [SKAT/README.md](SKAT/README.md) for the full list and usage.

## License

Apache License 2.0 — see [LICENSE](LICENSE).

## Citation

If you use this code or data, please cite the manuscript.

## Contact

- Zihao (John) Li — <>
- Christopher Fang-Yen (PI) — <>

For questions or issues, please contact the Fang-Yen Lab at The Ohio State University.
