## Supplementary material for "A semi-automated genetic screen using genome-wide association analysis identifies modulators of stress-induced sleep in *Caenorhabditis elegans*": Software S1: README.md

# SKAT analysis for stress-induced sleep (SIS) phenotype of MMP strains

A SKAT (Sequence Kernel Association Test) analysis was performed to quantify the association between individual genes in Million Mutation Project (MMP) *C. elegans* strains and a continuous stress-induced sleep (SIS) phenotype.

---

## 1. Overview

### Inputs to the pipeline

* The MMP variant data.
* The sleep phenotype for each strain (`combined_phenotype.csv`).

### Workflow

The pipeline filters the variants down to non-synonymous variants, groups them
by gene, matches each strain's genotype to its phenotype, and runs SKAT to compute the association with variation in SIS for every gene. See [Section 3](#3-running-the-pipeline) for how to run it.

### Outputs from the pipeline

* `SKAT_all-pvals.results` / `SKAT_all-qvals.results` — a p-value and q-value for every gene, indicating the strength of association to the phenotype.
* `SKAT_filtered_Na5.results` — the final ranked list used for candidate screening, limited to genes with enough alleles to test reliably (≥5 tested non-synonymous alleles).

---

## 2. File description

The SKAT folder contains three child folders: [scripts](scripts/), [inputs](inputs/), and [outputs](outputs/).

### Scripts

| File                    | What it does                                                                                               |
| ----------------------- | ---------------------------------------------------------------------------------------------------------- |
| `mmp_txt_to_vcf.py`     | Converts the MMP tabular download into `MMP.vcf` (see ["Building `MMP.vcf`"](#331-building-mmpvcf-from-the-mmp-database)). |
| `make_gene_variants.py` | Filters the raw MMP VCF down to [non-synonymous](#332-building-the-variant-list-gene_variantstxt) variants. |
| `run_pipeline.R`        | Builds the gene-to-variant map, attaches phenotypes, and runs SKAT.                                        |
| `Make_SSID_file.R`      | Standalone helper to build the gene-to-variant map from the command line.                                  |

The following scripts are kept in `scripts/debug/` for reference and are not part of the main pipeline:

| File            | What it does                                                     |
| --------------- | ---------------------------------------------------------------- |
| `SKAT_Run1.R`   | Older step-by-step version of the prep (SSID + phenotype setup). |
| `SKAT_prelim.R` | Interactive version of the SKAT run.                             |

### Inputs

The pre-generated input files are bundled in `inputs/inputs.zip`, which contains `gene_variants.txt`,
`MMP.bed`, `MMP.bim`, and `MMP.fam`. Unzip it into `inputs/` before running the pipeline
(`combined_phenotype.csv` is already unzipped in `inputs/`).

| File                        | What it is                                                                                                                                                                                              |
| --------------------------- | ------------------------------------------------------------------------------------------------------------------------------------------------------------------------------------------------------- |
| `MMP.vcf`                   | The raw Million Mutation Project variant data (not included due to large size ~6 Gb). Not required to run the pipeline with the pre-generated inputs (see [§3.2](#32-commands-necessary-to-run-the-pipeline-using-pre-generated-input-files)); only needed if rebuilding inputs from scratch (see [§3.3](#33-instructions-to-generate-input-files-from-scratch)). Built from the [MMP data portal](https://genome.sfu.ca/mmp/) download, see ["Building `MMP.vcf`"](#331-building-mmpvcf-from-the-mmp-database). |
| `gene_variants.txt`         | The filtered variant list, containing only the [non-synonymous](#332-building-the-variant-list-gene_variantstxt) variants.                                                                               |
| `MMP.bed` / `.bim` / `.fam` | Strain genotypes in PLINK format; the FAM file also holds the phenotype.                                                                                                                                   |
| `combined_phenotype.csv`    | The sleep phenotype Z-scores for each individually screened MMP strain.                                                                                                                                       |
| `MMP.SSID`                  | A map telling SKAT which variants belong to which gene (built during the run).                                                                                                                          |

### Outputs

| File                        | What it is                                                                                                                                  |
| --------------------------- | ------------------------------------------------------------------------------------------------------------------------------------------- |
| `SKAT_all-pvals.results`    | A p-value for every gene after Bonferroni correction, a lower p implies stronger association with changes in phenotype                     |
| `SKAT_all-qvals.results`    | The sorted version of SKAT_all-pvals.results with an additional column showing q-values after FDR control.                                  |
| `SKAT_filtered_Na5.results` | The ranked list limited to genes with at least 5 tested non-synonymous alleles (6,663 genes) — the final list used for candidate selection. |
| `validation_genes.tsv`      | The 15 known sleep genes used to validate the ranking, with their rank and percentile.                                                      |

---

## 3. Running the pipeline

### 3.1 Install required dependencies

#### R packages required

```r
install.packages(c("SKAT", "fdrtool", "dplyr", "plyr", "stringr"))
```

#### PLINK

Only needed if rebuilding `MMP.bed` from the VCF. Install with conda:

```bash
conda install -c bioconda plink
```

Or download the binary directly from [www.cog-genomics.org/plink](https://www.cog-genomics.org/plink/).


### 3.2 Commands necessary to run the pipeline using pre-generated input files

All the input files necessary to run the pipeline have been pre-generated and compressed in `inputs.zip`, so it just takes two commands to run the pipeline:

```bash
unzip inputs/inputs.zip -d inputs/
Rscript scripts/run_pipeline.R
```

`run_pipeline.R` does four things in order:

| Step | What it does                                                 | Produces                                           |
| ---- | ------------------------------------------------------------ | -------------------------------------------------- |
| 1    | Build the gene-to-variant map (defines 19,749 genes)         | `MMP.SSID`                                         |
| 2    | Match each strain's phenotype to its genotype                | updated `MMP.fam`                                  |
| 3    | Run SKAT on every gene, then add FDR q-values and sort       | `SKAT_all-pvals.results`, `SKAT_all-qvals.results` |
| 4    | Re-rank the genes that have ≥5 tested non-synonymous alleles | `SKAT_filtered_Na5.results` (final)                |

Temporary files (`MMP.SSID`, `MMP.SSD`, etc.) are written to `inputs/` and regenerated each run.

---

### 3.3 Instructions to generate input files from scratch

This section is only for users who want to generate input files from scratch.

---

#### 3.3.1 Building `MMP.vcf` from the MMP database

The MMP data portal (https://genome.sfu.ca/mmp/) distributes the variant calls as a tab-separated table, not as a VCF. Download "Results from mutagenized strains" (file size: 76 MB, if it comes as `.xlsx`, export the first sheet to a `.txt`/`.tsv`), then convert it to a VCF:

```bash
python3 scripts/mmp_txt_to_vcf.py --txt mmp_mut_strains_data.txt --out MMP.vcf
```

This produces the `MMP.vcf` (857,114 variants for 2,007 strains) used by the steps below. The INFO field is written with the `GF=`, `SN=`, `PN=`, `AAC=`, and `INDEL=` tags
that `make_gene_variants.py` reads.

---

#### 3.3.2 Building the variant list (`gene_variants.txt`)

We keep a variant only if it is non-synonymous:

* **SNVs** (single-nucleotide variants)
* **Indels** (insertions/deletions)
* **Structural variants**

Everything else is dropped: synonymous, intronic, intergenic, UTR, and non-coding-RNA variants. This leaves 191,938 variants (from ~857k in the raw VCF).

```bash
python3 scripts/make_gene_variants.py --vcf MMP.vcf --out inputs/gene_variants.txt \
        --verify inputs/gene_variants.txt   # optional: check against a reference
```

This reproduces the deposited `gene_variants.txt`.

> A plain `grep` for coding tags does not reproduce this file. It keeps the synonymous and
> intronic variants too. Use the script.
>
> This filter is also why some genes never appear in the results. For example, *flp-13* is
> represented by three variants, but all are in the non-coding region, so they're dropped here.

---

#### 3.3.3 Building the genotype files (`MMP.bed` / `.bim` / `.fam`)

To generate the PLINK files from the raw VCF, you need to build the strain-keep list, then convert:

```bash
awk -F',' 'NR>1 {print $1, $1}' inputs/combined_phenotype.csv > keep_strains.txt

plink --vcf MMP.vcf \
      --keep keep_strains.txt \
      --double-id \
      --allow-extra-chr \
      --allow-no-sex \
      --make-bed \
      --out inputs/MMP
```

> `--allow-extra-chr` is needed because *C. elegans* uses Roman-numeral chromosome names (I–V, X).
> Strains not in the VCF are dropped silently. The strain order in `MMP.fam` must match `MMP.bed`;
> `run_pipeline.R` keeps them aligned when it updates phenotypes, so don't reorder the FAM by hand.

---

## 4. Outputs

### 4.1 Some important gene counts

| Counts |  Meaning                                                                                                  |
| ------- | --------------------------------------------------------------------------------------------------------- |
| 19,749  | genes represented by ≥1 non-synonymous alleles in the entire MMP set (2,007 strains)                       |
| 18,070  | genes represented by ≥1 non-synonymous alleles in the 940 MMP strains we screened                         |
| 1,679   | genes not represented by any non-synonymous alleles in the 940 MMP strains we screened, left unranked (P = 1) |
| 6,663   | genes represented by ≥5 non-synonymous alleles in the 940 MMP strains we screened (the final ranked list for candidate screening)                                     |

See [§3.3.2](#332-building-the-variant-list-gene_variantstxt) for the definition of non-synonymous variants.

---

### 4.2 File and column descriptions

#### `SKAT_all-pvals.results`
Results for all 19,749 genes

| Column | Description |
|---|---|
| `SetID` | Gene sequence name (e.g. `ZK1067.1`) |
| `P.value` | SKAT P-value (1 if untested) |
| `N.Marker.All` | Variants assigned to the gene for the entire MMP set |
| `N.Marker.Test` | Variants present in the MMP strains we screened and actually used by SKAT (0 = not present) |

#### `SKAT_all-qvals.results`
Same as `SKAT_all-pvals.results`, but sorted by P-value, plus a column of FDR `Q.value` (`fdrtool`). The 1,679 untested genes (P = 1)
sort to the bottom.

#### `SKAT_filtered_Na5.results`
The ranked list for genes represented by at least 5 tested non-synonymous alleles (`N.Marker.Test >= 5`),
sorted by P-value. This list contains 6,663 genes and was used for candidate screening. We chose
the threshold of 5 by analyzing the median and mean percentile rankings of known sleep
genes across allele-count cutoffs (see main manuscript for details).

#### `validation_genes.tsv`
The 15 known sleep genes used to validate the ranking, with their rank and percentile among the
18,070 tested genes.

| Column | Description |
|---|---|
| `gene` | Gene name (e.g. `aptf-1`) |
| `SetID` | Gene sequence name (e.g. `K06A1.1`) |
| `P.value` | SKAT P-value |
| `N.Marker.All` | Variants assigned to the gene for the entire MMP set |
| `N.Marker.Test` | Variants present in the MMP strains we screened and actually used by SKAT |
| `rank_of_18070_tested` | Rank of this gene among the 18,070 tested genes (1 = strongest association) |
| `percentile` | Percentile rank of this gene among the 18,070 tested genes |
| `source` | Database the gene's `SetID` (gene sequence name) is sourced from (e.g. `WormBase`) |
