## Supplementary material for "A semi-automated genetic screen using genome-wide association analysis identifies modulators of stress-induced sleep in *Caenorhabditis elegans*": Software S1: SKAT_prelim.R

```
# load SKAT library (to do association analysis)
library(SKAT)
library(fdrtool)
library(dplyr)
library(plyr)


# Generate a SNP set data file (`.SSD`) and an `.info` file from binary plink formated data files using user specified SNP sets.
Generate_SSD_SetID('/Users/sreekarkutagulla/Desktop/data/MMP.bed', '/Users/sreekarkutagulla/Desktop/data/MMP.bim', '/Users/sreekarkutagulla/Desktop/data/MMP.fam', '/Users/sreekarkutagulla/Desktop/data/MMP.SSID', '/Users/sreekarkutagulla/Desktop/data/MMP.SSD', '/Users/sreekarkutagulla/Desktop/MMP.info')

# read in fam file
fam_file <- read.table(file = "/Users/sreekarkutagulla/Desktop/data/MMP.fam")

# open .SSD and .info files we just created
SSD.info <- Open_SSD('/Users/sreekarkutagulla/Desktop/data/MMP.SSD', '/Users/sreekarkutagulla/Desktop/MMP.info')

# create null model based on phenotypes (and co-variates if you have any)
set.seed(100)
Null_Model <- SKAT_Null_Model(fam_file$V6 ~ 1, out_type="C")

# perform SKAT on all sets of variants
#All_SKAT_Data  <- SKAT.SSD.All(SSD.info, Null_Model) 
All_SKAT_Data  <- SKAT.SSD.All(SSD.INFO = SSD.info, obj = Null_Model) 

head(All_SKAT_Data$results)

# sort SKAT results by p-value
head(All_SKAT_Data$results[order(All_SKAT_Data$results$P.value),])

#save SKAT results
write.table(x = All_SKAT_Data$results, file = "/Users/sreekarkutagulla/Desktop/data/SKAT_all-pvals.results", row.names = FALSE, col.names = TRUE, quote = FALSE, append = FALSE)

# calculate q-values
qvals <- fdrtool(All_SKAT_Data$results$P.value, statistic = "pvalue", cutoff.method="fndr", plot = FALSE)
All_SKAT_Data$results$Q.value <- qvals$qval

# sort SKAT results by q-value
All_SKAT_Data$results <-All_SKAT_Data$results[order(All_SKAT_Data$results$P.value),]
All_SKAT_Data$results[1:20,]

#save SKAT results
write.table(x = All_SKAT_Data$results, file = "/Users/sreekarkutagulla/Desktop/data/SKAT_all-qvals.results", row.names = FALSE, col.names = TRUE, quote = FALSE, append = FALSE)

SSID <- read.table(file = "/Users/sreekarkutagulla/Desktop/data/MMP.SSID", stringsAsFactors = FALSE)

library(plyr)

# count how many variants are in each snp set
#MACs <- count(SSID[,1])

# plot this as a histogram
MAC_hist <- hist(x <- (MACs[MACs$freq < 50,2]), breaks = 50, xlab = "Minor Allele Count", ylab = "Number of Genes", main = "Allele distribution")

# load .SSID file and give meaningful column names
SSID <- read.table(file = "/Users/sreekarkutagulla/Desktop/data/MMP.SSID", header=FALSE)
colnames(SSID) <- c("gene", "variant")
head(SSID)

# make a list of the variants to keep (i.e. plyr::count the number of occurrences of each gene)
#SSID_count <- count(SSID[,1])
head(SSID_count)

# make SSID file with only those variants (i.e. reduce the SSID list to only those with >= 6 variants)
genes_to_include <- data.frame(SSID_count[SSID_count$freq >= 4, 1])

# give the data frame meaningful column names  
colnames(genes_to_include) <- ("gene")
head(genes_to_include)

# reduce SSID to contain only those genes from the list genes_to_include via dplyr::semi_join
reduced_SSID <- semi_join(SSID, genes_to_include)
head(reduced_SSID)

# save SSID file
write.table(x = reduced_SSID, file = "/Users/sreekarkutagulla/Desktop/data/MMP-reduced.SSID", row.names = FALSE, col.names = FALSE, quote = FALSE)


Generate_SSD_SetID('/Users/sreekarkutagulla/Desktop/data/MMP.bed', '/Users/sreekarkutagulla/Desktop/data/MMP.bim', '/Users/sreekarkutagulla/Desktop/data/MMP.fam', '/Users/sreekarkutagulla/Desktop/data/MMP-reduced.SSID', '/Users/sreekarkutagulla/Desktop/data/MMP-reduced.SSD', '/Users/sreekarkutagulla/Desktop/MMP-reduced.info')

# open .SSD and .info files we just created
SSD.info_reduced <- Open_SSD('/Users/sreekarkutagulla/Desktop/data/MMP-reduced.SSD', '/Users/sreekarkutagulla/Desktop/MMP-reduced.info')

# null model doesn't take in info from .SSID, so we don't need to run that again
# perform SKAT on all sets of variants
All_SKAT_Data_reduced  <- SKAT.SSD.All(SSD.INFO = SSD.info_reduced, obj = Null_Model)

# sort SKAT results by p-value
head(All_SKAT_Data_reduced$results[order(All_SKAT_Data_reduced$results$P.value),])

# calculate q-values
qvals <- fdrtool(All_SKAT_Data_reduced$results$P.value, statistic = "pvalue", cutoff.method="fndr", plot = FALSE)
All_SKAT_Data_reduced$results$Q.value <- qvals$qval

# sort SKAT results by q-value
All_SKAT_Data_reduced$results <- All_SKAT_Data_reduced$results[order(All_SKAT_Data_reduced$results$P.value),]
All_SKAT_Data_reduced$results[1:20,]

#save SKAT results
write.table(x = All_SKAT_Data_reduced$results, file = "/Users/sreekarkutagulla/Desktop/data/SKAT_all_reduced.results", row.names = FALSE, col.names = TRUE, quote = FALSE, append = FALSE)
```
