## Supplementary material for "A semi-automated genetic screen using genome-wide association analysis identifies modulators of stress-induced sleep in *Caenorhabditis elegans*": Software S1: SKAT_Run1.R

```
library(dplyr)
library(stringr)


gene_variants <- read.table(file = "/Users/sreekarkutagulla/Desktop/data/gene_variants.txt", stringsAsFactors = FALSE)

pattern  <- "SN=[a-zA-Z0-9.]{1,}|CODING=[a-zA-Z0-9.]{1,}"
gene  <- str_extract(gene_variants$V8, pattern)
gene  <- sub("SN=|CODING=", "", gene)

SSID <- data.frame(gene, gene_variants$V3)
SSID <- subset(SSID, gene != "NA")
write.table(x = SSID, file = "/Users/sreekarkutagulla/Desktop/data/MMP.SSID", row.names = FALSE, col.names = FALSE, append = FALSE, quote = FALSE)

pheno_file <- read.csv(file = "/Users/sreekarkutagulla/Desktop/data/combined_phenotype.csv", header = TRUE, stringsAsFactors = FALSE)

# Read fam keeping all 6 PLINK columns (FID IID PID MID SEX PHENO)
fam_file <- read.table(file = "/Users/sreekarkutagulla/Desktop/data/MMP.fam", stringsAsFactors = FALSE)
colnames(fam_file) <- c("Strain", "IID", "PID", "MID", "SEX", "PHENO")

# left_join preserves fam strain order (critical: must match MMP.bed sample order)
fam_w_phenos <- left_join(fam_file, pheno_file, by = "Strain")
fam_w_phenos$PHENO <- fam_w_phenos$Phenotype
fam_w_phenos$Phenotype <- NULL

write.table(x = fam_w_phenos, file = "/Users/sreekarkutagulla/Desktop/data/MMP.fam", row.names = FALSE, col.names = FALSE, quote = FALSE, append = FALSE)
```
