## Supplementary material for "A semi-automated genetic screen using genome-wide association analysis identifies modulators of stress-induced sleep in *Caenorhabditis elegans*": Software S1: make_gene_variants.py

```
#!/usr/bin/env python3
"""
make_gene_variants.py  --  Build inputs/gene_variants.txt from the raw MMP VCF.

This is the FILTERING step that turns the full Million Mutation Project VCF
(~6.4 GB, all strains, all variant classes) into the coding, protein-altering
variant list used by the SKAT pipeline. It reproduces the deposited
gene_variants.txt exactly (byte-identical variant set: 191,938 variants).

WHAT IT KEEPS (a variant passes if ANY of the following holds):
  1. SNV  -- nonsynonymous coding change: INFO has AAC=X->Y with X != Y
            (missense or nonsense/stop). Synonymous (X->X) and non-coding
            SNVs (AAC=NA) are dropped.
  2. INDEL -- any induced small indel: INFO contains "INDEL=".
  3. SV   -- structural variant (deletion/etc.) carrying a coding annotation:
            INFO contains "SVTYPE=" AND "CODING=".
  AND, for SNVs and INDELs, the variant must map to a real gene: INFO has
  SN=<gene> where <gene> is not a chromosome name (I,II,III,IV,V,X,MtDNA),
  not empty/NA, and not a non-standard model (E_..., cTel...).

WHAT IT DROPS: intergenic, intronic, synonymous, UTR-only, ncRNA, and any
  variant with no real gene assignment.

OUTPUT: the 8 fixed VCF columns only (CHROM POS ID REF ALT QUAL FILTER INFO);
  the ~2007 per-strain genotype columns are discarded. No strain subsetting
  is performed -- every strain's variants are represented.

USAGE:
  python3 scripts/make_gene_variants.py --vcf MMP.vcf --out inputs/gene_variants.txt
  python3 scripts/make_gene_variants.py --vcf MMP.vcf --out inputs/gene_variants.txt \
          --verify inputs/gene_variants.txt    # check byte-for-byte against a reference

Downstream: run_pipeline.R reads inputs/gene_variants.txt to build the SKAT
  SSID (gene -> variant map). See scripts/run_pipeline.R and README.md.
"""

import argparse
import re
import sys

CHROMS = {"I", "II", "III", "IV", "V", "X", "MtDNA"}
_SN = re.compile(r"SN=([^;]*)")
_AAC = re.compile(r"AAC=([^;]*)")


def real_gene(info):
    """Return the gene name if INFO maps to a real gene, else None."""
    m = _SN.search(info)
    if not m:
        return None
    sn = m.group(1)
    if sn in CHROMS or sn in ("", "NA") or sn.startswith("E_") or sn.startswith("cTel"):
        return None
    return sn


def is_nonsynonymous(info):
    """True if AAC=X->Y with X != Y (missense or stop)."""
    m = _AAC.search(info)
    if not m or "->" not in m.group(1):
        return False
    aac = m.group(1)
    left, right = aac.split("->")[0], aac.split("->")[-1]
    return left != right


def keep(info):
    """Apply the full coding / protein-altering filter to one INFO field."""
    if "INDEL=" in info:                       # induced small indels
        return real_gene(info) is not None
    if "SVTYPE=" in info:                       # structural variants
        return "CODING=" in info
    # SNVs: nonsynonymous coding change on a real gene
    return is_nonsynonymous(info) and (real_gene(info) is not None)


def build(vcf_path, out_path):
    n_in = n_out = 0
    with open(vcf_path) as fin, open(out_path, "w") as fout:
        for line in fin:
            if line[0] == "#":
                continue
            n_in += 1
            # split off only the first 8 columns; INFO is column 8 (index 7)
            fields = line.split("\t", 8)
            info = fields[7]
            if keep(info):
                fout.write("\t".join(fields[:8]) + "\n")
                n_out += 1
    return n_in, n_out


def verify(out_path, ref_path):
    """Compare the variant-ID set of out_path against a reference file."""
    def ids(path):
        s = set()
        with open(path) as fh:
            for line in fh:
                s.add(line.split("\t")[2])
        return s
    a, b = ids(out_path), ids(ref_path)
    if a == b:
        print(f"VERIFY OK: {len(a)} variants match {ref_path} exactly.")
        return True
    print(f"VERIFY FAILED: produced {len(a)}, reference {len(b)}; "
          f"missing {len(b - a)}, extra {len(a - b)}.", file=sys.stderr)
    return False


def main():
    ap = argparse.ArgumentParser(description=__doc__,
                                 formatter_class=argparse.RawDescriptionHelpFormatter)
    ap.add_argument("--vcf", required=True, help="path to the raw MMP.vcf")
    ap.add_argument("--out", required=True, help="output gene_variants.txt path")
    ap.add_argument("--verify", metavar="REF",
                    help="after building, verify the ID set matches this reference file")
    args = ap.parse_args()

    n_in, n_out = build(args.vcf, args.out)
    print(f"Read {n_in} variant records; wrote {n_out} to {args.out}")

    if args.verify:
        if not verify(args.out, args.verify):
            sys.exit(1)


if __name__ == "__main__":
    main()
```
