## Supplementary material for "A semi-automated genetic screen using genome-wide association analysis identifies modulators of stress-induced sleep in *Caenorhabditis elegans*": Software S1: Make_SSID_file.R

```
##Make snp(variant) set ID (SSID) file for SKAT

main <- function() {
  
  ## Start the clock!
  ptm <- proc.time() 
  
  ## load libraries
  library(stringr)
  
  ## assign command line arguments
  args <- commandArgs(trailingOnly = TRUE)
  path_to_vcf_wout_header <- args[1]
  SSID.file.name <- args[2]
  
  ## make SSID file for my set of MMP strains
  ## note - takes some time to read in MMPdyf_non-syn_coding_no_header.txt (derived from merged .vcf file)
  my.SSID <- make_SSID(path_to_vcf_wout_header)
  
  ## save SSID file
  write.table(my.SSID, SSID.file.name, sep="\t", row.names=FALSE, col.names=FALSE, quote=FALSE, append=FALSE)
  
  ## output time to run script
  the_time <- proc.time() - ptm # Stop the clock
  print(paste("It took", the_time[3], "to run Make_SSID_file.R")) # Print time to load data
}

## Make snp(variant) set ID (SSID) file for SKAT
make_SSID <- function(vcfwoutheader.filename) {
  ## takes merged .vcf file and creates a dataframe from which you can save a snp(variant) set ID (SSID) 
  ## file for SKAT analysis
  
  ## import vcf file without header
  mydata.vcf <- read.table(vcfwoutheader.filename)
  
  ## get variant names
  variants <- mydata.vcf$V3
  
  ## get sequence names
  seq_tag  <- "SN=[a-zA-Z0-9.]{1,}|CODING=[a-zA-Z0-9.]{1,}"
  sequence_names  <- str_extract(mydata.vcf$V8, seq_tag)
  sequence_names  <- sub("SN=|CODING=", "", sequence_names)
  
  vars <- as.character(variants)
  seqs  <- as.character(sequence_names)
  
  SSID  <- data.frame(cbind(seqs, vars))
  return(SSID)
}

main()
```
