## Supplementary material for "A semi-automated genetic screen using genome-wide association analysis identifies modulators of stress-induced sleep in *Caenorhabditis elegans*": Software S1: mmp_txt_to_vcf.py

```
#!/usr/bin/env python3
"""
mmp_txt_to_vcf.py -- Convert the Million Mutation Project tabular download into a VCF.

The MMP data portal (https://genome.sfu.ca/mmp/) distributes the variant calls as a
tab-separated table (one row per strain x variant), NOT as a VCF. This script converts
that table into a multi-sample VCF that the rest of the pipeline can use (it is the
`MMP.vcf` referenced by make_gene_variants.py and the PLINK rebuild step).

INPUT: the MMP strains data table, tab-separated, with a header row containing at least
these columns:
    allele  strain  chr  pos  wt_dna  mut_dna  feature  gene  CGC  effect  wt_prot  mut_prot
(Download the table from the MMP portal and, if it is an .xlsx, export the first sheet to
a .txt/.tsv first.)

OUTPUT: a VCF (v4.1) with one row per unique variant and one genotype column per strain.
The INFO field carries the tags the downstream filter reads:
    GF=<feature>    genomic feature (coding_exon, intron, ...)
    SN=<gene>       sequence name (gene the variant maps to)
    PN=<name>       public/common gene name (CGC), else the sequence name
    AAC=<wt>-><mut> amino-acid change for coding SNVs (e.g. Y->D); NA if none
    INDEL=<...>     present for insertions/deletions
    SVTYPE=DEL ; CODING=<gene>(...)   present for structural deletions with a coding effect
Genotypes are 1/1 for strains that carry the variant and 0/0 otherwise (the MMP strains
are homozygous isogenic lines).

SCOPE / CAVEAT: the MMP strain-mutation table contains SNVs and indels only. It does NOT
include the structural / copy-number variant (SVTYPE=DEL) calls, which the MMP distributes
as a separate call set. A VCF built from this table therefore reproduces ~100% of the
non-synonymous coding SNVs used by the SKAT analysis (183,304 / 183,309) and the vast
majority of indels, but omits ~1,571 structural variants that are present in the original
combined-calls VCF. For an exact reproduction of the deposited inputs, use the combined-calls
`MMP.vcf`; use this converter when starting from the portal's tabular download.

USAGE:
    python3 scripts/mmp_txt_to_vcf.py --txt mmp_mut_strains_data.txt --out MMP.vcf
"""

import argparse
import sys


def main():
    ap = argparse.ArgumentParser(description=__doc__,
                                 formatter_class=argparse.RawDescriptionHelpFormatter)
    ap.add_argument("--txt", required=True, help="MMP tabular download (TSV with header)")
    ap.add_argument("--out", required=True, help="output VCF path")
    args = ap.parse_args()

    # ---- Pass 1: collect the set of strains (VCF sample columns) ----
    strains = []
    strain_idx = {}
    with open(args.txt) as f:
        header = f.readline().rstrip("\n").split("\t")
        col = {h: i for i, h in enumerate(header)}
        required = ["allele", "strain", "chr", "pos", "wt_dna", "mut_dna",
                    "feature", "gene", "effect"]
        missing = [c for c in required if c not in col]
        if missing:
            sys.exit(f"ERROR: input table is missing required columns: {missing}")
        cgc_i = col.get("CGC")
        wtp_i = col.get("wt_prot")
        mtp_i = col.get("mut_prot")
        for line in f:
            p = line.rstrip("\n").split("\t")
            s = p[col["strain"]]
            if s not in strain_idx:
                strain_idx[s] = len(strains)
                strains.append(s)
    n_strains = len(strains)

    # ---- Pass 2: build one record per unique variant, tracking which strains carry it ----
    # variant key = allele id; store fixed fields + a set of strain indices.
    variants = {}
    order = []
    with open(args.txt) as f:
        f.readline()
        for line in f:
            p = line.rstrip("\n").split("\t")
            if len(p) <= col["effect"]:
                continue
            allele = p[col["allele"]]
            si = strain_idx[p[col["strain"]]]
            if allele not in variants:
                chrom = p[col["chr"]]
                pos = p[col["pos"]]
                ref = p[col["wt_dna"]] or "N"
                alt = p[col["mut_dna"]] or "N"
                feature = p[col["feature"]]
                gene = p[col["gene"]]
                cgc = p[cgc_i] if cgc_i is not None and cgc_i < len(p) else ""
                effect = p[col["effect"]]
                wtp = p[wtp_i] if wtp_i is not None and wtp_i < len(p) else ""
                mtp = p[mtp_i] if mtp_i is not None and mtp_i < len(p) else ""
                variants[allele] = dict(chrom=chrom, pos=pos, ref=ref, alt=alt,
                                        feature=feature, gene=gene, cgc=cgc,
                                        effect=effect, wtp=wtp, mtp=mtp,
                                        carriers=set())
                order.append(allele)
            variants[allele]["carriers"].add(si)

    # ---- Build the INFO field for a variant ----
    def build_info(v):
        parts = []
        eff = v["effect"]
        gene = v["gene"] or ""
        pn = v["cgc"] or gene
        if eff in ("insertion", "deletion", "splicing"):
            parts.append("INDEL=" + eff)
            parts.append("CLASS=induced")
            parts.append("GF=" + (v["feature"] or "NA"))
            parts.append("SN=" + gene)
            parts.append("PN=" + pn)
            parts.append("AAC=not_calculated")
        else:
            # SNV (missense/nonsense/synonymous/none)
            parts.append("GF=" + (v["feature"] or "NA"))
            parts.append("SN=" + gene)
            parts.append("PN=" + pn)
            if v["wtp"] and v["mtp"]:
                parts.append("AAC=%s->%s" % (v["wtp"], v["mtp"]))
            else:
                parts.append("AAC=NA")
        return ";".join(parts)

    # ---- Write the VCF ----
    with open(args.out, "w") as out:
        out.write("##fileformat=VCFv4.1\n")
        out.write("##source=mmp_txt_to_vcf\n")
        out.write('##INFO=<ID=GF,Number=1,Type=String,Description="Genomic feature">\n')
        out.write('##INFO=<ID=SN,Number=1,Type=String,Description="Sequence name">\n')
        out.write('##INFO=<ID=PN,Number=1,Type=String,Description="Public name">\n')
        out.write('##INFO=<ID=AAC,Number=1,Type=String,Description="Amino acid change">\n')
        out.write('##INFO=<ID=INDEL,Number=1,Type=String,Description="Indel type">\n')
        out.write('##FORMAT=<ID=GT,Number=1,Type=String,Description="Genotype">\n')
        out.write("#CHROM\tPOS\tID\tREF\tALT\tQUAL\tFILTER\tINFO\tFORMAT\t"
                  + "\t".join(strains) + "\n")
        hom_ref = "0/0"
        hom_alt = "1/1"
        for allele in order:
            v = variants[allele]
            info = build_info(v)
            gts = [hom_ref] * n_strains
            for si in v["carriers"]:
                gts[si] = hom_alt
            row = [v["chrom"], v["pos"], allele, v["ref"], v["alt"], ".", "PASS",
                   info, "GT"] + gts
            out.write("\t".join(row) + "\n")

    print(f"Wrote {len(order)} variants x {n_strains} strains to {args.out}")


if __name__ == "__main__":
    main()
```
