## Supplementary material for "A semi-automated genetic screen using genome-wide association analysis identifies modulators of stress-induced sleep in *Caenorhabditis elegans*": Software S1: run_pipeline.R

```
library(plyr)
library(dplyr)
library(stringr)
library(SKAT)
library(fdrtool)

# Run this script from the SKAT_MMP/ root directory: Rscript scripts/run_pipeline.R
data_dir <- "inputs"
out_dir  <- "outputs"

# ── Step 1: Generate SSID ────────────────────────────────────────────────────
cat("Generating SSID...\n")
gene_variants <- read.table(file.path(data_dir, "gene_variants.txt"), stringsAsFactors = FALSE)
pattern <- "SN=[a-zA-Z0-9.]{1,}|CODING=[a-zA-Z0-9.]{1,}"
gene <- str_extract(gene_variants$V8, pattern)
gene <- sub("SN=|CODING=", "", gene)
SSID <- subset(data.frame(gene, gene_variants$V3), gene != "NA")
write.table(SSID, file.path(data_dir, "MMP.SSID"),
            row.names = FALSE, col.names = FALSE, append = FALSE, quote = FALSE)

# ── Step 2: Update MMP.fam phenotypes ───────────────────────────────────────
cat("Updating MMP.fam phenotypes...\n")
pheno_file <- read.csv(file.path(data_dir, "combined_phenotype.csv"),
                       header = TRUE, stringsAsFactors = FALSE)
fam_file <- read.table(file.path(data_dir, "MMP.fam"), stringsAsFactors = FALSE)
colnames(fam_file) <- c("Strain", "IID", "PID", "MID", "SEX", "PHENO")
fam_w_phenos <- left_join(fam_file, pheno_file, by = "Strain")
fam_w_phenos$PHENO <- fam_w_phenos$Phenotype
fam_w_phenos$Phenotype <- NULL
write.table(fam_w_phenos, file.path(data_dir, "MMP.fam"),
            row.names = FALSE, col.names = FALSE, quote = FALSE, append = FALSE)

# ── Step 3: Full SKAT run ────────────────────────────────────────────────────
cat("Generating SSD (full)...\n")
ssd_full  <- file.path(data_dir, "MMP.SSD")
info_full <- file.path(data_dir, "MMP.info")
if (file.exists(ssd_full))  file.remove(ssd_full)
if (file.exists(info_full)) file.remove(info_full)
Generate_SSD_SetID(
  file.path(data_dir, "MMP.bed"),
  file.path(data_dir, "MMP.bim"),
  file.path(data_dir, "MMP.fam"),
  file.path(data_dir, "MMP.SSID"),
  ssd_full, info_full
)

fam_final   <- read.table(file.path(data_dir, "MMP.fam"))
SSD.info    <- Open_SSD(ssd_full, info_full)
set.seed(100)
Null_Model  <- SKAT_Null_Model(fam_final$V6 ~ 1, out_type = "C")

cat("Running SKAT (full)...\n")
All_SKAT_Data <- SKAT.SSD.All(SSD.INFO = SSD.info, obj = Null_Model)

write.table(All_SKAT_Data$results,
            file.path(out_dir, "SKAT_all-pvals.results"),
            row.names = FALSE, col.names = TRUE, quote = FALSE)

qvals <- fdrtool(All_SKAT_Data$results$P.value,
                 statistic = "pvalue", cutoff.method = "fndr", plot = FALSE)
All_SKAT_Data$results$Q.value <- qvals$qval
All_SKAT_Data$results <- All_SKAT_Data$results[order(All_SKAT_Data$results$P.value), ]
write.table(All_SKAT_Data$results,
            file.path(out_dir, "SKAT_all-qvals.results"),
            row.names = FALSE, col.names = TRUE, quote = FALSE)

# ── Step 4: Final filtered list (genes with >= 5 tested non-synonymous alleles) ──
# N.Marker.Test is the number of alleles SKAT actually tested for each gene
# (after dropping markers with MAF = 0 or excess missingness). We keep genes with
# at least 5 tested alleles, re-rank them by p-value, and write the final list.
cat("Filtering to genes with >= 5 tested alleles (N.Marker.Test)...\n")
filtered <- All_SKAT_Data$results[All_SKAT_Data$results$N.Marker.Test >= 5, ]
filtered <- filtered[order(filtered$P.value), ]
cat("Genes with >= 5 tested alleles:", nrow(filtered), "\n")

write.table(filtered,
            file.path(out_dir, "SKAT_filtered_Na5.results"),
            row.names = FALSE, col.names = TRUE, quote = FALSE)

cat("Done. Results saved to", out_dir, "\n")
cat("Top 10 genes (final filtered list):\n")
print(head(filtered[, c("SetID", "P.value", "Q.value")], 10))
```
