## Supplementary material for "A semi-automated genetic screen using genome-wide association analysis identifies modulators of stress-induced sleep in *Caenorhabditis elegans*": Software S1: Make_Table_S1.m

```
%% Combine Fig2 and Fig3 supplemental CSVs into one workbook (like Table_S1.xlsx).
% Sheet "MMP_screen"       <- figure2/Fig2_MMP_quiescence_data.csv
% Sheet "candidate_screen" <- figure3/Fig3_mutants_quiescence_data.csv
%
% Run this script from anywhere; paths below are resolved relative to
% this file's location (tableS1/), one level under the repo root.

clear; clc;

repoRoot = fileparts(fileparts(mfilename('fullpath')));

fig2Csv = fullfile(repoRoot, 'figure2', 'Fig2_MMP_quiescence_data.csv');
fig3Csv = fullfile(repoRoot, 'figure3', 'Fig3_mutants_quiescence_data.csv');
outFile = fullfile(fileparts(mfilename('fullpath')), 'Table_S1.xlsx');

if isfile(outFile)
    delete(outFile); % avoid stale sheets from a previous run
end

mmpScreen = readtable(fig2Csv);

% Force SKAT_Rank_Percentile to import as text: readtable auto-detects it
% as numeric (since most rows are a single number) and would otherwise
% convert both "NA" and multi-locus values like "45.1; 32.97" to NaN.
fig3Opts = detectImportOptions(fig3Csv);
fig3Opts = setvartype(fig3Opts, 'SKAT_Rank_Percentile', 'string');
candidateScreen = readtable(fig3Csv, fig3Opts);

writetable(mmpScreen, outFile, 'Sheet', 'MMP_screen');
writetable(candidateScreen, outFile, 'Sheet', 'candidate_screen');

fprintf('Wrote %s\n', outFile);
fprintf('  MMP_screen: %d rows\n', height(mmpScreen));
fprintf('  candidate_screen: %d rows\n', height(candidateScreen));
```
